# Spatiotemporal control of cell cycle acceleration during axolotl spinal cord regeneration

**DOI:** 10.1101/2020.02.10.941443

**Authors:** Emanuel Cura Costa, Leo Otsuki, Aida Rodrigo Albors, Elly M. Tanaka, Osvaldo Chara

**Author notes:** These authors contributed equally to this work (Emanuel Cura Costa, Leo Otsuki). Corresponding author: Osvaldo Chara, Center for Information Services and High Performance Computing (ZIH), Technische Universität Dresden, Nöthnitzer Straße 46, 01187 Dresden, Germany., Systems Biology Group (SysBio), Institute of Physics of Liquids and Biological Systems (IFLySIB), National Scientific and Technical Research Council (CONICET) and University of La Plata, Calle 59 N 789, 1900 La Plata, Argentina., Web: http://sysbioiflysib.wordpress.com/.

## Abstract

Axolotls are uniquely able to resolve spinal cord injuries, but little is known about the mechanisms underlying spinal cord regeneration. We previously found that tail amputation leads to reactivation of a developmental-like program in spinal cord ependymal cells (Rodrigo Albors *et al*., 2015), characterized by a high-proliferation zone emerging 4 days post-amputation (Rost *et al*., 2016). What underlies this spatiotemporal pattern of cell proliferation, however, remained unknown. Here, we use modelling, tightly linked to experimental data, to demonstrate that this regenerative response is consistent with a signal that recruits ependymal cells during 85 hours after amputation within ^~^830□m of the injury. We adapted FUCCI technology to axolotls (AxFUCCI) to visualize cell cycles in vivo. AxFUCCI axolotls confirmed the predicted appearance time and size of the injury-induced recruitment zone and revealed cell cycle synchrony between ependymal cells. Our modeling and imaging move us closer to understanding bona fide spinal cord regeneration.

## Introduction

The axolotl (*Ambystoma mexicanum*) has the remarkable ability to regenerate the injured spinal cord (reviewed in Freitas, Yandulskaya & Monaghan, 2019; Tazaki *et al*, 2017; Chernoff *et al*., 2003), and thus represents a unique system to study the mechanisms of successful spinal cord regeneration. Key players in this process are the ependymal cells lining the central canal of the spinal cord, which retain neural stem cell potential throughout life (Becker, Becker & Hugnot, 2018).

In earlier studies, we found that spinal cord injury in the axolotl triggers the reactivation of a developmental-like program in ependymal cells, including a switch from slow, neurogenic to fast, proliferative cell divisions (Rodrigo Albors *et al*., 2015). We showed that in the uninjured spinal cord and in the non-regenerating region of the injured spinal cord, ependymal cells divide slowly, completing a cell cycle in 14.2 ± 1.3 days. In contrast, regenerating ependymal cells speed up their cell cycle and divide every 4.9 ± 0.4 days (Rodrigo Albors *et al*., 2015; Rost et al., 2016). By using a mathematical modeling approach, we demonstrated that the acceleration of the cell cycle is the major driver of regenerative spinal cord outgrowth and that other processes such as cell influx, cell rearrangements, and neural stem cell activation play smaller roles (Rost *et al*., 2016). We quantitatively analyzed cell proliferation in space and time and identified a high-proliferation zone that emerges 4 days after amputation within the 800 μm adjacent to the injury site and shifts posteriorly over time as the regenerating spinal cord grows (Rost *et al*., 2016)(Figure 1–figure supplement 1). What underlies this precise spatiotemporal pattern of cell proliferation in the regenerating axolotl spinal cord, however, remains unknown. Pattern formation phenomena occurring during development can be quantitatively reproduced by invoking morphogenetic signals spreading from localized sources (Morelli *et al*., 2012). It is thus conceivable that tail amputation triggers a signal that propagates or diffuses along the injured spinal cord to speed up the cell cycle of resident cells.

In this new study, we take a modelling approach supported by previous and new experimental data to unveil the spatiotemporal distribution that such a signal should have in order to explain the observed rate of spinal cord outgrowth in the axolotl. We generate a new transgenic axolotl that reports cell cycle phases to confirm several of our theoretical predictions *in vivo*. Together, our results provide new clues for when and where to search for the signal/s that may be responsible for driving successful spinal cord regeneration.

## Results

### Model of uninjured spinal cord

Taking into account the symmetry of the ependymal tube and that ependymal cells organize as a pseudo-stratified epithelium (Joven & Simon, 2018), we modeled the anterior-posterior (AP) axis of the spinal cord as a row of ependymal cells (see Computational methods section 1.1 for more details). We modeled ependymal cells as rigid spheres of uniform diameter and assumed that they can be either cycling or quiescent and defined the fraction of cycling cells as the growth fraction, *GF*. We modeled the proliferation dynamics of cycling cells as follows: we assumed that in the initial condition, each cycling cell is in a random coordinate along its cell cycle, where the initial cell cycle coordinate and the cell cycle length follow an exponential and a lognormal distribution, respectively. In the uninjured axolotl spinal cord, upon cell division, i) the daughter cells inherit the cell cycle length from the mother’s lognormal distribution and ii) the daughter cells translocate posteriorly, displacing the cells posterior to them. This last feature of the model is the implementation of what we earlier defined as “cell pushing mechanism” (Rost *et al*., 2016). This model predicts that after a time of approximately one cell cycle length, mitotic events will occur along the AP axis and contribute to the growth of the spinal cord (Figure 1A).

**Figure 1.**
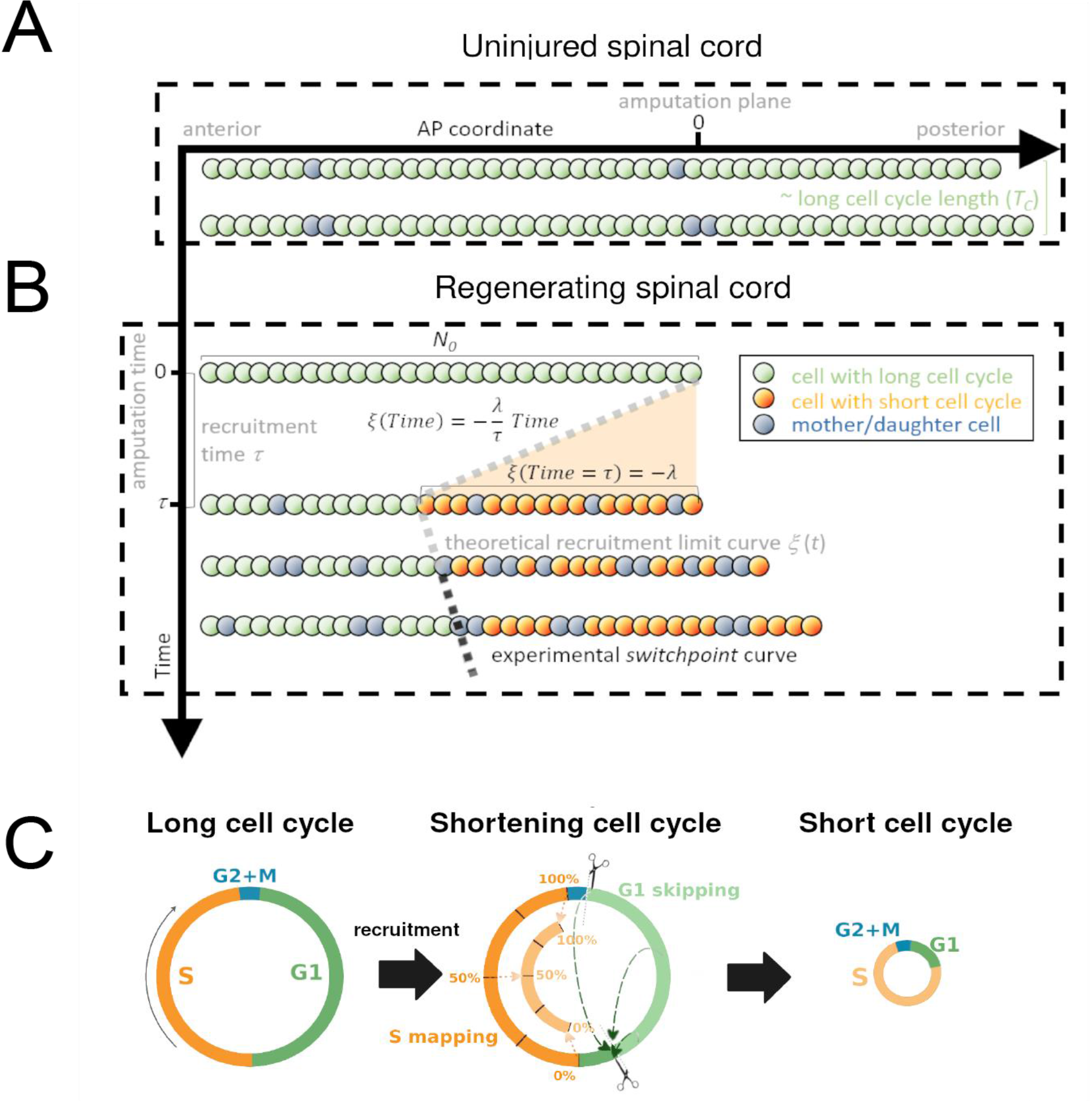
Model of uninjured and regenerating spinal cord growth based on shortening G1 and S cell phases. **A) Uninjured spinal cord**. 1D model of ependymal cells aligned along the AP axis. In uninjured tissue, ependymal cells cycle with long cell cycle length (see Figure 1 – figure supplement 1) and when they divide “push” cells posteriorly and the spinal cord tissue grows. As an example, two mother cells (in blue) in the top row give rise to four daughter cells in the second row (also in blue) following one division. This results in a growth of two cell diameters within a timeframe of approximately one long cell cycle length. **B) Regenerating spinal cord.** After amputation (AP coordinate and time equal to zero) a signal is released anteriorly from the amputation plane during a time *τ* and spreads while recruiting resident ependymal cells up to the theoretical recruitment limit *ξ* located at – *λ* μm from the amputation plane. After a certain time, the recruitment limit *ξ* overlaps the experimental switchpoint (see Figure 1 – figure supplement 1). **C) Proposed mechanisms of cell cycle shortening consists of partial skipping of G1 phase and proportional mapping between long and short S phases.** In the middle panel, two examples are depicted (dashed green arrows) of cells which were in the long G1 phase (immediately before recruitment) that become synchronized (immediately after recruitment). Additionally, there are three examples (dotted orange arrows) of cells that were in the long S phase (immediately before recruitment) that are proportionally mapped (immediately after recruitment). All these examples are shown in detail in Figure 1 – figure supplement 2. The diameter of the circles is approximately proportional to the length of the cell’s cell cycle.

### Model of regenerating spinal cord

Next, we removed the most posterior cells of the tissue to model tail amputation and the regenerative response in the remaining cells (*N_0_*) *in silico* (see Figure 1B and Computational methods section 1.1 for more details). We assumed that amputation triggers the release of a signal that spreads anteriorly from the injury site with constant speed along the AP axis, recruiting cells by inducing a change in their cell cycle. We established that cell recruitment stops at time *τ*, recruiting *λ* μm of cells anterior to the amputation plane. We notated the AP position of the most anterior cell recruited by the signal as *ξ*(*t*), and called this the recruitment limit, such that *ξ*(*t* = *τ*) = −*λ*. In the model, all cycling cells anterior to the cell located at *ξ*(*t*) are not recruited and continue cycling slowly during the simulations (Figure 1–figure supplement 2). In contrast, cycling cells posterior to *ξ*(*t*) are recruited at a time *t* within the interval 0 ≤ *t* ≤ *τ* and irreversibly modify their cycling according to their cell cycle phase at the time of recruitment.

Because we previously demonstrated that the length of G2 and M phases do not change upon amputation (Rodrigo Albors *et al*., 2015), we assumed that cells in G2 or M within the recruitment zone will continue cycling as before (Figure 1C, Computational methods section 1.1.3). In contrast, because we showed that regenerating cells go through shorter G1 and S phases than non-regenerative cells (G1 shortens from 152 ± 54 hours to 22 ± 19 hours; S shortens from 179 ± 21 hours to 88 ± 9 hours, Rodrigo Albors *et al*., 2015), we reasoned that the signal instructs recruited cells to shorten G1 and S, effectively shortening their cell cycle.

To explain how ependymal cells may shorten G1 and S phase in response to the injury signal, we conceived a mechanism of G1 shortening in which a certain part of this cell cycle phase is skipped. We implemented this mechanism as follows (Figure 1C, Figure 1 – figure supplement 2, Computational methods section 1.1.1): if at the time of recruitment the cell is in G1, there are two possible coordinate transformations. If the cell cycle coordinate is located before the difference between the long (non-regenerative) G1 length and the shortened (regenerative) G1 length, the cell clock is reset; *i.e*., its transformed cell cycle coordinate will be the beginning of the shortened G1 phase in the next simulation step (immediately after G2 + M phases). In contrast, if the original cell cycle coordinate of the cell is located after the difference between the long G1 length and the shortened G1 length, there is no change in the time to enter into S phase. The difference between the long and short G1 lengths constitutes a sort of threshold. If the G1 cell lies before this value, it skips and if it lies after, it continues cycling as before. This mechanism of G1 skipping predicts a partial synchronization of the cell cycle as cells transit through G1 (Figure 1 – figure supplement 2) – an important implication that we test experimentally later.

Because all DNA must be duplicated prior to cell division, we considered a different mechanism to model S phase shortening: if the cell cycle coordinate belongs to S at the moment of recruitment, the new cell cycle coordinate of this cell will be proportionally mapped to the corresponding coordinate of a reduced S phase in the next simulation step (Figure 1C, Figure 1 – figure supplement 2, Computational methods section 1.1.2). For instance, if the recruited cell is 40% into its (long) S phase when the signal arrives, it will be in the 40% of its shorter S phase in the next simulation step.

Daughter cells of recruited cells inherit short G1 and S phases from their mothers and consequently have shorter cell cycle lengths (Figure 1C). Finally, we assumed that recruitment of a quiescent cell would induce its progress from G0 to G1 after an arbitrary delay (Computational methods section 1.1.4). To parametrize the cell phase durations of recruited and non-recruited cycling cells, the growth fraction and cell geometry, we used our previous experimental data from regenerating and non-regenerating regions of axolotl spinal cords (Rodrigo Albors *et al*., 2015)(Computational methods section 1.2 and Table 1).

**Table 1.**
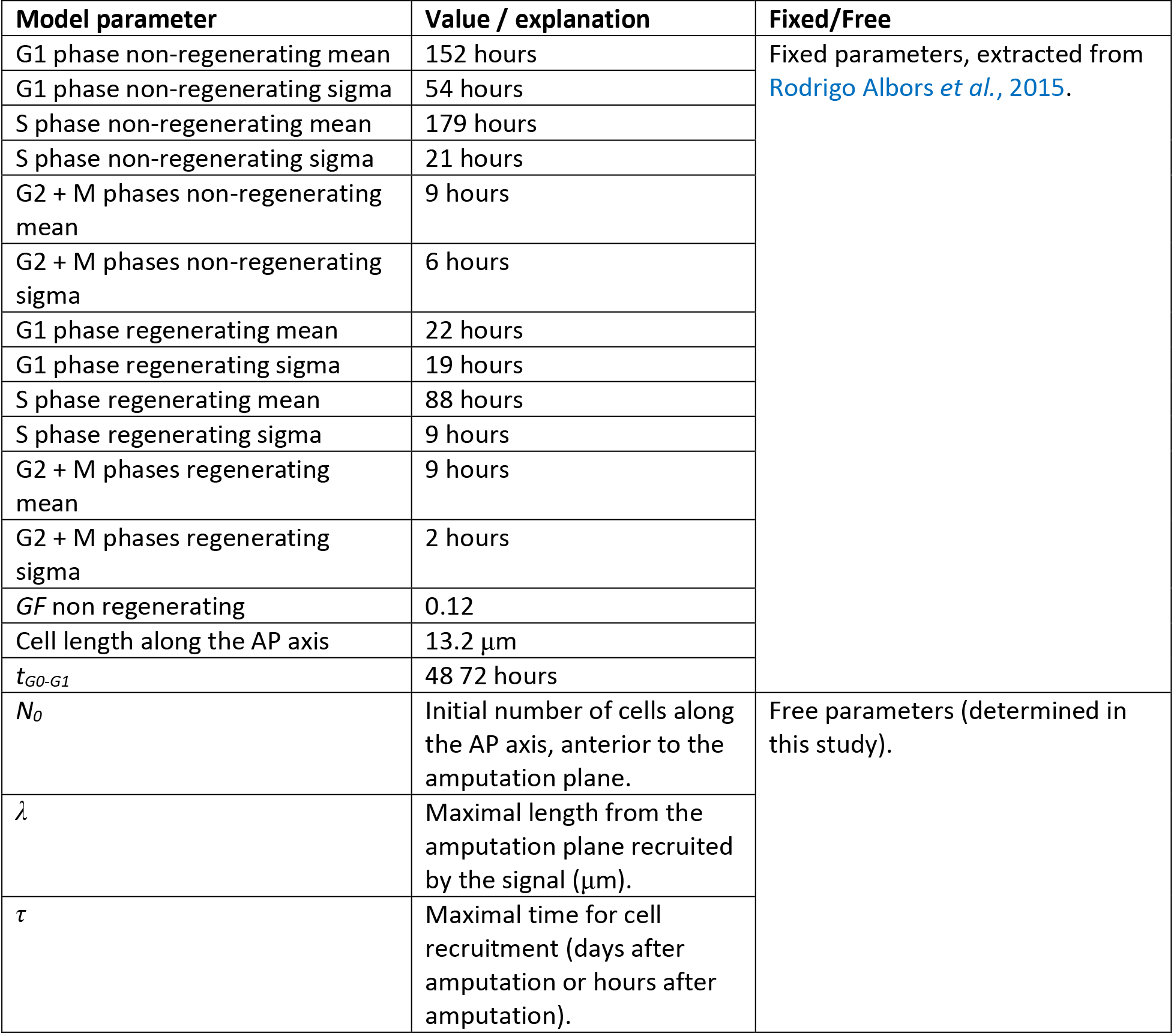
Model parameters.

The model predicts that if we wait a time similar to the short cell cycle length, we will observe more proliferation posterior to *ξ* than anterior to it. In particular, if this model is correct, the prediction for *ξ*(Figure 1B) should agree with the experimental curve of the switchpoint separating the region of high proliferation from low proliferation along the AP axis of the axolotl spinal cord during regeneration (Figure 1-figure supplement 1).

### Regenerative spinal cord outgrowth can be explained by a signal that acts during the 85 hours following amputation and recruits cells within 828 μm of the amputation plane

To evaluate if the model could explain the regenerative outgrowth of the ependymal tube and to estimate the free parameters, we fitted *ξ*(*t*) to the experimental switchpoint (Rost *et al*., 2016). Specifically, we followed an inference Approximate Bayesian Computation method, streamlined by the use of pyABC-framework (Klinger, Rickert & Hasenauer, 2018). The model successfully reproduced the experimental switchpoint with best fitting parameters *N_0_* = 196 ± 2 cells, *λ* = 828 ± 30 μm and *τ* = 85 ± 12 hours (Figure 2A; parameter posterior distributions obtained after convergence are shown in Figure 2-figure supplement 1 and see computational methods section 1.3 for details). Interestingly, a clonal analysis of the model shows that while the anterior-most cells are slightly displaced, cells located close to the amputation plane end up at the posterior end of the regenerated spinal cord (Figure 2–figure supplement 2A), in agreement with cell trajectories observed during axolotl spinal cord regeneration (Rost *et al*., 2016). Additionally, the velocity of a clone monotonically increases with its AP coordinate (Figure 2–figure supplement 2B), also in line with experimental data (Rost *et al*., 2016). These results suggest that cells preserve their relative position along the AP axis. Hence, when plotting the relative position of each clone to the outgrowth of the corresponding tissue minus the recruitment limit *ξ*(*t*), we observed that this normalized quantity is conserved in time, a fingerprint of the scaling behavior characteristic of regeneration (Figure 2–figure supplement 2C). Importantly, with the parameterization leading to the best fitting of the experimental switchpoint, we quantitatively predicted the time evolution of the regenerative outgrowth that was observed *in vivo* (Rost *et al*., 2016) (Figure 2B, Video 1).

**Figure 2.**
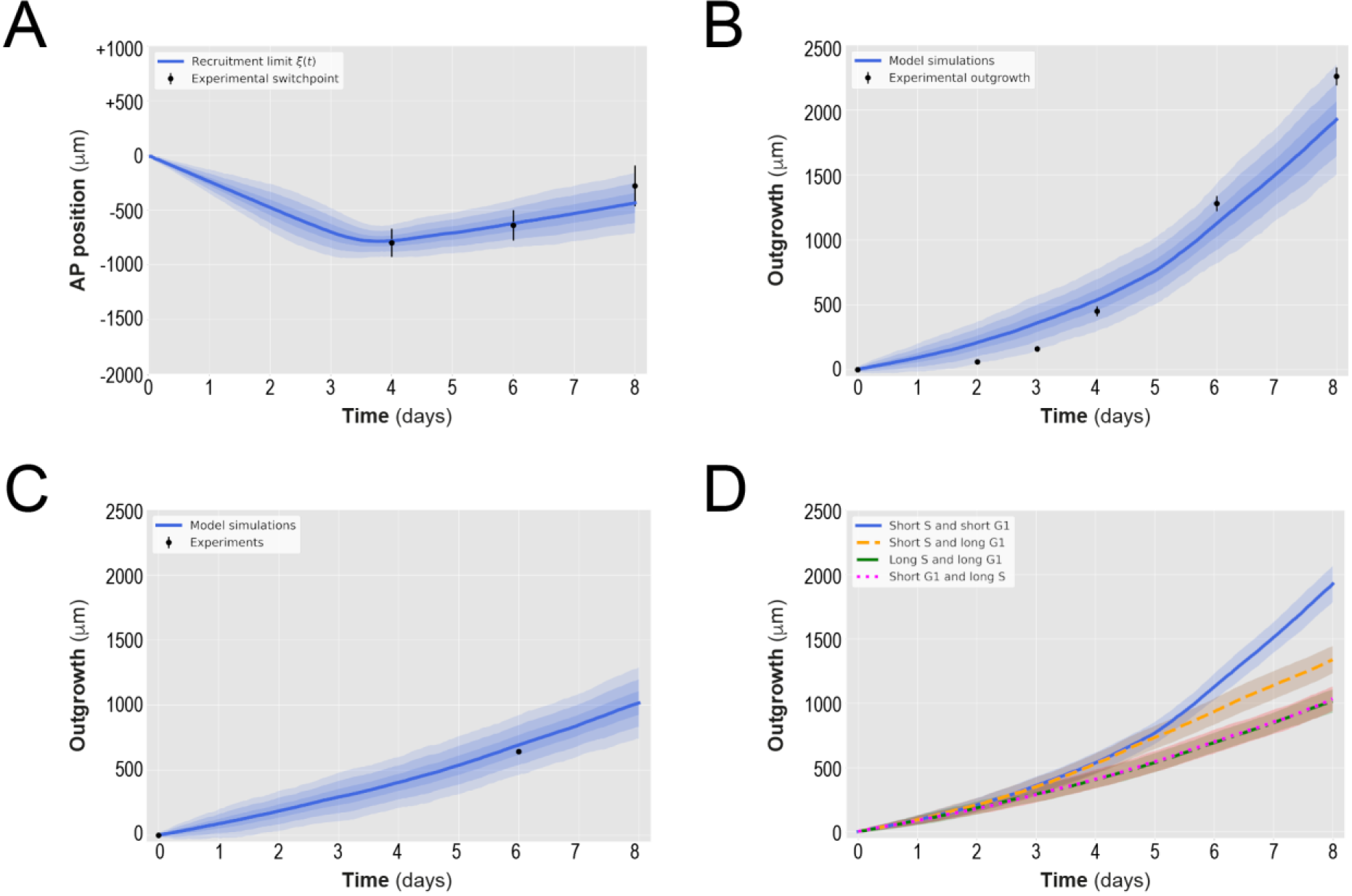
A hypothetical signal recruits ependymal cells up to 828 μm anterior to the amputation plane during the 85 hours following amputation, recapitulating *in vivo*_spinal cord regenerative outgrowth. **A) The modeled recruitment limit successfully fits the experimental switchpoint curve**. Best fitting simulations of the model-predicted recruitment limit *ξ*(*t*) overlap the experimental switchpoint curve. Best fitting parameters are *N_0_* = 196 ± 2 cells, *λ* = 828 ± 30 μm and *τ* = 85 ± 12 hours. **B) The model quantitatively matches experimental axolotl spinal cord outgrowth kinetics** (Rost *et al*., 2016). **C) The model reproduces experimental outgrowth reduction when the acceleration of cell proliferation is impeded.** Prediction of the model assuming that neither S nor G1 phase lengths were shortened superimposed with experimental outgrowth kinetics in which acceleration of the cell cycle was prevented by knocking out *Sox2* (Fei *et al*., 2014). Lines in A, B and C show the means while blue shaded areas correspond to 68, 95 and 99.7 % confidence intervals, from darker to lighter, calculated from the 1,000 best fitting simulations. **D) S phase shortening dominates cell cycle acceleration during axolotl spinal cord regeneration**. Spinal cord outgrowth kinetics predicted by the model assuming shortening of S and G1 phases (blue line), only shortening of S phase (orange dashed line), only shortening of G1 phase (magenta dotted line) and neither S nor G1 shortening (green line). The magenta line and green line are overlapped with one another. Means are represented as lines and each shaded area corresponds to one sigma out of 1,000 simulations.

Our model assumed that cells are rigid spheres of uniform diameter fixed from the mean length of ependymal cells measured along the AP axis (Rost *et al*., 2016). To test whether this *naïve* assumption could impact on our prediction of the regenerative spinal cord outgrowth, we repeated the simulations but replacing the mean cell length by the biggest and smallest possible cell lengths within a 99 % confidence interval based on earlier data (Rost *et al*., 2016). Spinal cord outgrowth predicted under these two extreme scenarios could hardly be distinguished from the previous prediction (Figure 2–figure supplement 3A, B). Similar results were obtained when we assumed that ependymal cells do not have a constant length along the AP axis but one extracted from a normal distribution parametrized from the experimental data on ependymal cell lengths along the AP axis (Figure 2–figure supplement 3C).

When we simulated a fast recruitment process by reducing the maximal recruitment time *τ* to one day while maintaining the maximal recruitment length *λ* constant, we found that the model-predicted outgrowth overestimates the experimental outgrowth (Figure 2–figure supplement 4A). On the contrary, when we decreased recruitment speed by increasing *τ* to 8 days while keeping *λ* constant, we observed a shorter outgrowth than that observed experimentally (Figure 2–figure supplement 4A). Reducing the maximal recruitment distance *λ* to zero mimics a hypothetical case in which the signal is incapable of recruiting the cells anterior to the amputation plane (Figure 2-figure supplement 4B). Increasing *λ* by approximately 100 % without changing *τ* (that is, increasing recruitment speed) results in more recruited ependymal cells and faster spinal cord outgrowth than observed *in vivo* (Figure 2–figure supplement 4B). These results point to a spatially and temporarily precise cell recruitment mechanism underlying the tissue growth response during axolotl spinal cord regeneration.

### Regenerating spinal cord outgrowth when cell cycle acceleration is impeded

We next asked how much the spinal cord would grow if the cell cycle acceleration instructed by the injury signal is blocked. We made use of our model and predicted the tissue outgrowth when the lengths of G1 and S were unaltered after amputation. In this condition, all cells would divide with the durations of cell cycle phases reported under non-regenerating conditions (Rodrigo Albors *et al*., 2015). Our results show that blocking recruitment, and therefore the acceleration of the cell cycle, slows down tissue growth, leading to an outgrowth of 694 ± 77 μm instead of the observed 1,127 ± 103 μm at day 6 (Figure 2C). This result is consistent with reducing down to zero the maximal recruitment length *λ* (Figure 2-figure supplement 4B). Interestingly, this model-predicted outgrowth is in agreement with the reported experimental outgrowth in *Sox2* knock-out axolotls, in which the acceleration of the cell cycle does not take place after amputation (Figure 2C, Fei *et al*., 2014).

### S phase shortening is sufficient to explain the initial regenerative spinal cord outgrowth

The relative contributions of G1 *vs* S phase shortening to spinal cord outgrowth are an important unknown that is technically difficult to interrogate *in vivo*. We made use of our model to answer this question *in silico*. For this, we maintained the same parametrization recapitulating spinal cord outgrowth (Figure 2A and B) but modified the model such that recruited cycling cells shorten S phase but not G1 phase (*i.e*., leaving unaltered G1 phase) or *vice versa*. Interestingly, our results indicate that shortening of only S phase can explain the explosive spinal cord outgrowth observed *in vivo*, independently of G1 shortening, up to day 4 (Figure 2D, blue line and orange line). In contrast, shortening of only G1 phase has a mild impact on the initial outgrowth, as it results in an outgrowth almost identical to the case in which neither G1 nor S phase were reduced (Figure 2D, magenta line *versus* green line). From day 4, though, shortening of only S phase cannot recapitulate the observed outgrowth (Figure 2D, blue line and orange line) and indeed, it is the shortening of both S and G1 phases that returns the same outgrowth than that observed *in vivo*. These modeling predictions are a consequence of i) the proximity of S phase to the next cell division compared with G1 phase ii) the fact that S phase represents ~ 7.5 days of the total ~ 14 days of the long cell cycle, which is reduced to ~3.7 days in the ~5 days short cell cycle and iii) the time window of the investigated outgrowth was 8 days. To conclude, these results indicate that, up to day 4, shortening of S phase can explain the regenerative spinal cord outgrowth in the axolotl, while the effect of G1 shortening manifests from day 5.

### Visualizing cell cycle progression in axolotls *in vivo* using FUCCI

Our model makes defined assumptions on how the phases of the cell cycle shorten to result in an acceleration of cell cycle over 85 hours within 828 μm of the injury site. We sought to validate the model by determining the kinetics of this response rigorously using a tool that distinguishes cell cycle phases *in vivo* while preserving spatiotemporal context. For this, we adapted Fluorescent Ubiquitination-based Cell Cycle Indicator (FUCCI) technology to axolotls (Figure 3A) (Zielke & Edgar, 2015). FUCCI is a genetically encoded reporter that distinguishes cell cycle phases by capitalizing on the mutually exclusive, oscillatory activity of two ubiquitin ligases (Sakaue-Sawano *et al*., 2008). SCF^Skp2^ is active in S and G2 of the cell cycle, when it targets the DNA licensing factor Cdt1 for proteolytic degradation. In contrast, APC/C^Cdh1^ is active from mid-M to G1; during these phases, it targets Geminin (Gmnn; a Cdt1 inhibitor) for degradation. Fusing the degradation-targeting motifs (degrons) in the Cdt1 and Gmnn proteins to two distinct fluorophores puts fluorophore abundance under the control of SCF^Skp2^ and APC/C^Cdh1^ activity and enables fluorescence to be used as a readout for cell cycle phase. Importantly, analyzing FUCCI does not require cell dissociation (thus preserving spatial context within the tissue), immunostaining or measurement of DNA content.

**Figure 3.**
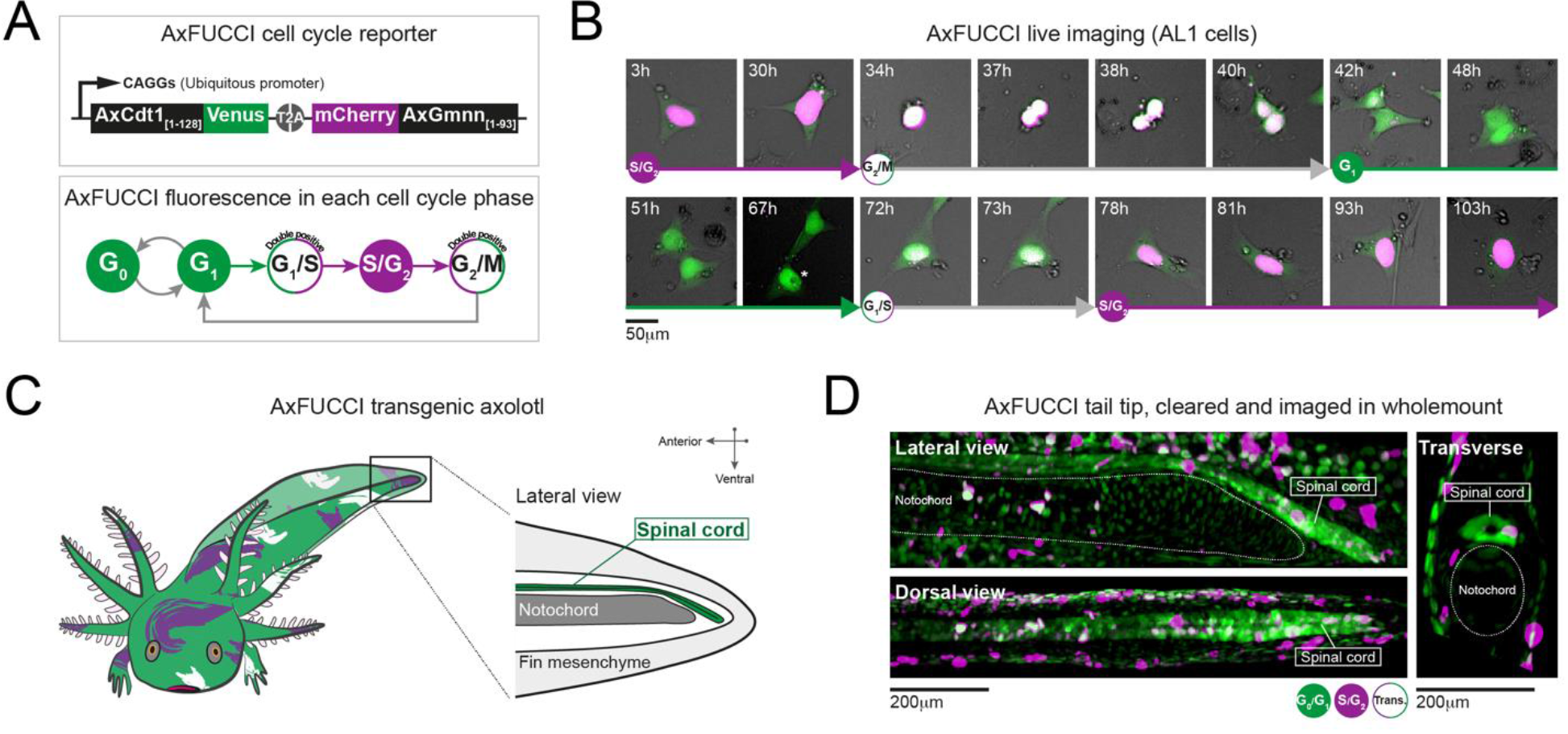
AxFUCCI – a transgenic cell cycle reporter for axolotl. **A) Top panel: AxFUCCI design.** A ubiquitous promoter (CAGGs) drives expression of two AxFUCCI probes (AxCdt1[1-128]-Venus and mCherry-AxGmnn[1-93]) in one transcript. The two AxFUCCI probes are separated co-translationally by virtue of the viral ‘self-cleaving’ T2A peptide sequence. **Bottom panel: AxFUCCI fluorescence combinations in each phase of the cell cycle. B) Live imaging of a single AxFUCCI-electroporated axolotl cell *in vitro*.** Each panel is a single frame acquired at the indicated hour (h) after the start of the imaging session. One cell cycle (from S/G2 to the subsequent S/G2) is depicted. At 67h, the two mitotic daughter cells moved apart; the asterisked daughter cell is depicted in the remaining panels. **C) Establishment of AxFUCCI transgenic axolotls.** The location of the spinal cord is indicated in the context of the tail. **D) A fixed AxFUCCI tail, cleared and imaged in wholemount using lightsheet microscopy.** The 3D data enable *post-hoc* digital sectioning of the same spinal cord into lateral, dorsal or transverse views. Images depict maximum intensity projections through 50 μm (lateral and transverse views) or 150 μm (dorsal view) of tissue. Trans.: Transition-AxFUCCI (G1/S or G2/M transition).

FUCCI has been adapted successfully to several model organisms, including mouse, zebrafish, *Drosophila* and human cells (reviewed by Zielke & Edgar, 2015). We designed axolotl FUCCI *de novo* by extracting the degron-harboring sequences from the axolotl Cdt1 and axolotl Gmnn proteins. This was important as we found that the N-terminus of Cdt1 protein, harboring the PIP degron, is divergent across animal models (Figure 3-figure supplement 1A). We defined the relevant fragments of axolotl Cdt1 protein (harboring the PIP degron and Cy motif) and axolotl Gmnn protein (harboring the D box degron) using homology alignment and comparison with zebrafish FUCCI (Figure 3-figure supplements 1A,B,C,D) (Sugiyama *et al*., 2009, Bouldin *et al*., 2014). The axolotl Cdt1_[aa1-128]_ fragment was fused to mVenus fluorescent protein and the axolotl Gmnn_[aa1-93]_ fragment to mCherry fluorescent protein. We used the CAGGs ubiquitous promoter and viral T2A sequence to co-express the Cdt1_[aa1-128]_-mVenus and Gmnn_[aa1-93]_-mCherry fusions in one transcript. The resulting axolotl-specific cell cycle reporter is referred to as AxFUCCI (Figure 3A).

### AxFUCCI discriminates cell cycle phases faithfully *in vivo*

We performed live cell imaging, DNA content analysis and immunofluorescence-based characterizations to validate the ability of AxFUCCI to report the phases of the cell cycle. First, we electroporated an immortalized axolotl cell culture line (AL1 cells) with AxFUCCI plasmid and performed live imaging. As expected, AxFUCCI fluorescence oscillated during the cell cycle in the order mVenus > Double positive > mCherry > Double positive > mVenus (Figure 3B, Figure 3-figure supplements 2A and B, Videos 2 and 3). We confirmed that mVenus-positive cells were G0/G1 cells and that mCherry-positive cells were S/G2 cells using a flow cytometer to analyze DNA content (Figure 3-figure supplement 3A). Cells are transiently AxFUCCI double positive between G0/G1 and S/G2 (Figure 3B and Figure 3-figure supplements 2A,B); we infer that these cells are at the G1/S boundary, as observed in mouse FUCCI (Abe *et al*. 2013). Interestingly, and in contrast to FUCCI in other model organisms, we observe a second AxFUCCI double positive window between S/G2 and G0/G1, corresponding to the G2/M boundary and M cells (Figure 3B and Figure 3-figure supplements 2A,B). Thus, AxFUCCI discriminates the following cell cycle phases in axolotl: G0/G1 (mVenus only); S/G2 (mCherry only); G1/S transition and G2/M transition (double positive) (Figure 3B). Importantly, AxFUCCI’s capacity to label defined landmarks in the cell cycle (*i.e*. G1/S and G2/M transition) enabled us to later test for cell cycle synchronization, a characteristic feature of our model.

We generated stable transgenic AxFUCCI axolotls using I-SceI-mediated transgenesis, bred them to sexual maturity and used F_1_ (germline-transmitted) progeny for further validations (Figure 3C). AxFUCCI animals developed at a similar rate to their non-transgenic siblings, did not differ in their basal cell proliferation (Figure 3-figure supplement 4A) and regenerated amputated tail tissue with similar kinetics to the *d/d* animals used in our previous study (Figure 3-figure supplement 4B). Cells dissociated from AxFUCCI axolotl tails and analyzed using a flow cytometer exhibited the expected fluorescence/DNA content relationships (Figure 3-figure supplement 4C). As a second assay, we prepared spinal cord tissue sections from AxFUCCI axolotls and compared DNA content (as assessed by DAPI fluorescence) in mVenus *vs* mCherry-positive cells. As expected, mCherry-positive cells harbored significantly more DNA than mVenus-positive cells (Figure 3-figure supplement 4D,E). Thirdly, we injected AxFUCCI axolotls intraperitoneally with EdU, a thymidine analogue that is incorporated into DNA during S phase. Following an 8 hour EdU pulse, mCherry-positive cells but not mVenus-positive cells should be EdU-positive and this was indeed the case (Figure 3-figure supplement 4F). Finally, we co-stained AxFUCCI spinal cord tissue sections with cell type-specific markers. NeuN-expressing neurons, located on the periphery of the spinal cord, are post-mitotic, differentiated cells (G0) and should be mVenus-positive (and never mCherry-positive). Indeed, we found that 100% of neurons that expressed AxFUCCI were mVenus-positive (Figure 3-figure supplement 4G). By contrast, Sox2-expressing ependymal cells, which are proliferative cells, expressed mVenus, mCherry or both fluorophores (Figure 3-figure supplement 4H).

Based on these validations, and for simplicity, we refer to mVenus fluorescence as ‘G0/G1-AxFUCCI’ (green in Figures 3-6), mCherry fluorescence as ‘S/G2-AxFUCCI’ (magenta in Figures 3-6) and double positivity as ‘Transition-AxFUCCI’ (white in Figures 3-6).

### Measuring the recruitment zone *in vivo* using AxFUCCI

We used AxFUCCI animals to measure the size of the ependymal cell recruitment zone *in vivo*. We amputated AxFUCCI tails at 5 mm from the tail tips and harvested replicate regenerating tails daily up to 5 days post-amputation. We implemented a pipeline to optically clear and image fixed AxFUCCI tail tissue in wholemount, which enabled *post-hoc* digital re-sectioning of the imaging data into any orientation for accurate measurement (Figure 3D and Video 4) (Pende & Vadiwala *et al*., 2020). Importantly, this pipeline did not alter the length of the tail tissue (Figure 3-figure supplement 5). We found that, at amputation, most ependymal cells expressed G0/G1-AxFUCCI and only a minority expressed S/G2-AxFUCCI (Figure 4A). In the five days following tail tip amputation, the proportion of S/G2-AxFUCCI-expressing cells increased locally and sharply at the amputation site, then propagated anteriorly along the spinal cord, consistent with the appearance of a recruitment zone (Figure 4A and Figure 1B).

**Figure 4.**
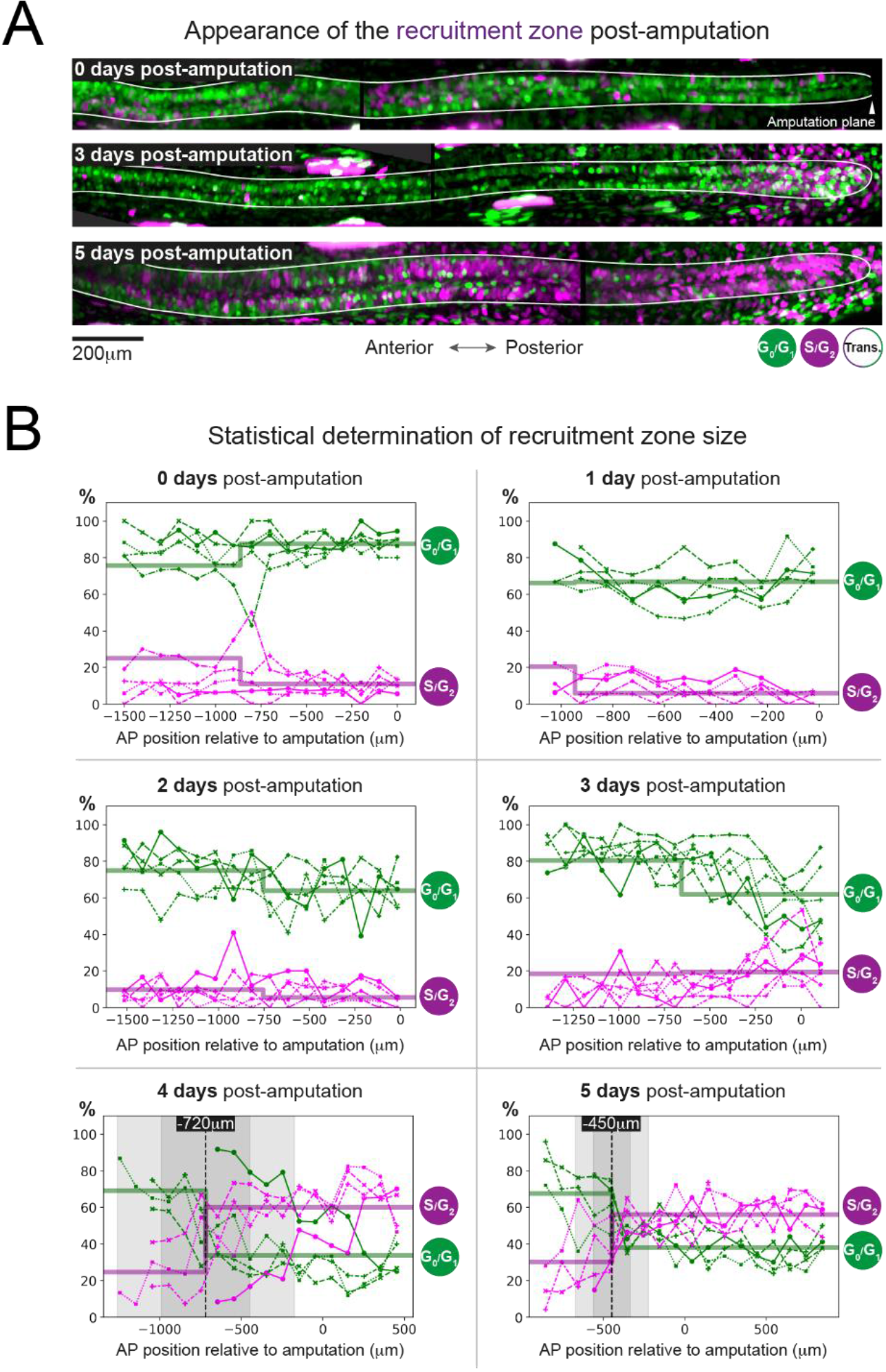
The predicted recruitment zone size is observed in AxFUCCI tails after amputation. **A) Visualization of the recruitment zone in AxFUCCI tails.** At amputation, most ependymal cells are G0/G1-AxFUCCI positive (green). After amputation, a decrease in G0/G1-AxFUCCI and an increase in S/G2-AxFUCCI (magenta) expression is seen anterior to the amputation plane and this zone increases in size anteriorly through day 5 post-amputation. The spinal cord is outlined. Images depict maximum intensity projections through 30 μm of tissue and are composites of two adjacent fields of view. **B) Quantification of the AP border that delimits the recruitment zone in G0/G1 and S/G2-AxFUCCI data.** Percentage of G0/G1 (in green) and S/G2 (in magenta)-AxFUCCI-expressing cells quantified in 100 μm bins along the anterior-posterior spinal cord axis. A mathematical model assuming two adjacent spatially homogeneous zones separated by an anterior-posterior border (AP border) was fitted to the G0/G1 and S/G2-AxFUCCI-data for each animal. Significant differences between anterior and posterior zones were detected only at day 4 and 5 (Kolmogorov-Smirnov test *p* = 0.0286). The AP border mean and two sigmas are depicted as a black dashed line and a grey shaded areas respectively. AP position is defined with respect to the amputation plane (0 μm). *n* = 4-6 tails per time point, ~ 300 cells each. Different symbols depict different animals; each line represents one animal. Best fitting values of the model regarding the anterior and posterior percentage of AxFUCCI data is in Figure 4 – figure supplement 1. Individual fittings are in Figure 4 – figure supplement 2. For more details, see Materials and Methods section 2.12.

As a first step, we quantified the percentage of ependymal cells expressing either G0/G1-AxFUCCI alone or S/G2-AxFUCCI alone within the 1000-1600 μm of spinal cord anterior to the injury site at each day post-amputation. A 1600 μm length should encompass not only the recruitment zone but also more anteriorly located ependymal cells that are not recruited by the injury signal and that continue to cycle slowly (Figure 1B). We performed our quantifications in 100 μm adjacent bins to preserve spatial information, using the severed notochord tip to denote the amputation plane. To test statistically whether the cells expressing G0/G1-AxFUCCI and S/G2-AxFUCCI are heterogeneously distributed along the AP axis in the regenerating spinal cords (*i.e*. if a recruitment zone can be detected), we followed an approach similar to that that we used previously to determine the switchpoint (Rost *et al*., 2016). We fitted the measured spatial AP profiles of the percentage of G0/G1-AxFUCCI and S/G2-AxFUCCI-expressing cells in each animal with a mathematical model assuming two adjacent homogeneous spatial zones separated by an anterior-posterior border (AP border), which we assumed was the same for both G0/G1 and S/G2-AxFUCCI data. For each animal at each time point, we tested if the mean percentage of cells expressing G0/G1-AxFUCCI or S/G2-AxFUCCI in the anterior *versus* the posterior zones were significantly different by running a Kolmogorov-Smirnov test. Up until 3 days post-amputation, no statistical significance was detected between anterior and posterior in the G0/G1-AxFUCCI and S/G2-AxFUCCI data (Figure 4B, Figure 4 -figure supplement 1A-D and Figure 4 -figure supplement 2A-D). By contrast, at 4 and 5 days post-amputation, G0/G1-AxFUCCI and S/G2-AxFUCCI data revealed a significant difference between the anterior and posterior zones, consistent with the appearance of a recruitment zone (Figure 4B, Figure 4 -figure supplement 1E, F and Figure 4 – figure supplement 2E,F).

Crucially, we measured the AP border of the AxFUCCI data to be at −717 ± 272 μm relative to the amputation plane on day 4 and −446 ± 112 μm on day 5, overlapping within two sigma the −782 ± 50 μm and −710 ± 62 μm recruitment limits predicted by our model (Figure 4B). Moreover, the appearance of the recruitment zone between days 3 and 4 post-amputation accommodates the 85 hour recruitment time in our model. Thus, AxFUCCI animals confirmed the predicted appearance time and size of the recruitment zone.

### Regenerating cells have high cell cycle synchrony *in vivo*

Our model of G1 shortening (Figure 1C and Figure 1-figure supplement 2) predicts that ependymal cells in the recruitment zone should exhibit high synchrony with each other in the cell cycle during early regeneration, a property that has not been investigated. AxFUCCI axolotls enabled us to assess for potential ependymal cell cycle synchrony in regenerating spinal cord.

We performed a more rigorous quantification of cell cycle distribution from our wholemount data in which we focused on the 600 μm of spinal cord immediately anterior to the amputation plane, within the recruitment zone, and also including Transition-AxFUCCI (G1/S and G2/M transition cells) and M phase cells. At the moment of amputation, 85 ± 5 % of ependymal cells expressed G0/G1-AxFUCCI and 11 ± 6 % expressed S/G2-AxFUCCI (Figure 5A). We note that this baseline differs from the one that we reported previously in smaller, *d/d* control animals, but we found this lower basal proliferation rate to be consistent among animals in this study independent of genotype (Figure 3-figure supplement 4A) (Rodrigo Albors *et al*., 2015). Restrictions in feeding and/or changes in animal handling during the COVID-19 pandemic might explain this difference (see Discussion). As we expected our model might be robust to the baseline cell cycle profile, we gathered measurements in these animals for further analysis.

**Figure 5.**
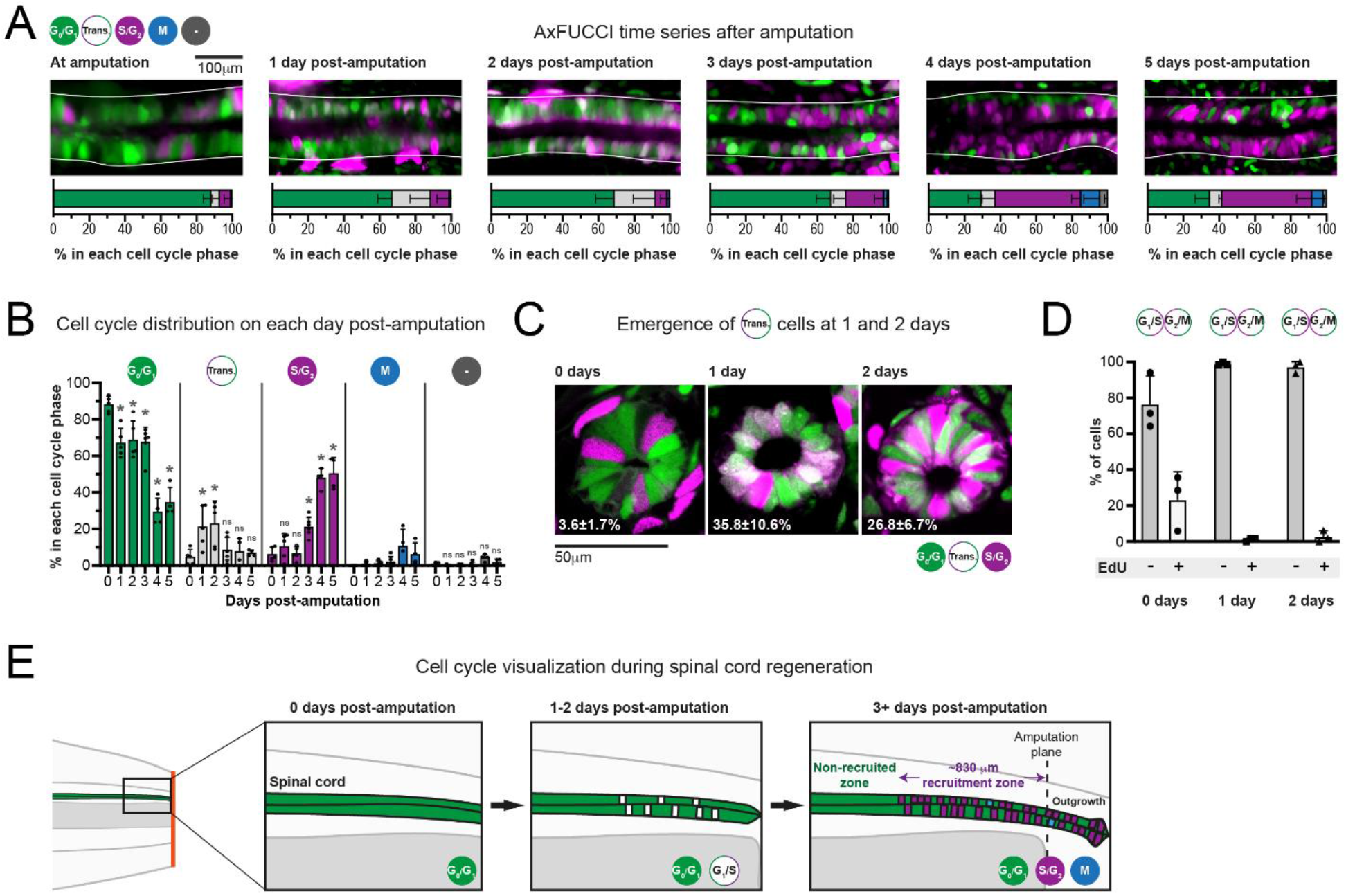
Ependymal cells exhibit cell cycle synchrony during spinal cord regeneration. **A) Cell cycle distributions of ependymal cells during the first five days of regeneration.** Top row: maximum intensity projections through 25 μm of spinal cord oriented anterior to left and posterior to right. The spinal cord is outlined. The lumen is in the center. Images were taken from within the 600 μm of most posterior regenerating spinal cord. Bottom row: percentage of ependymal cells in each cell cycle phase in the 600 μm of most posterior regenerating spinal cord. Cells in mitosis (M) were counted independently of AxFUCCI and were identified based on their condensed chromatin as revealed by staining with DAPI (not shown). ‘-’ indicates AxFUCCI-negative cells, which were negligible at all time points. *n* = 4-6 tails per time point, ~ 100 cells each. Error bars indicate standard deviations. **B) AxFUCCI data separated by cell cycle phase.** Quantitative data from panel A re-plotted by cell cycle phase. Day 0 data (immediately after amputation) were taken as baselines, and statistical analyses were performed against these baselines. Kruskal-Wallis tests followed by Wilcoxon rank sum tests revealed a significant decrease in G0/G1-AxFUCCI from 1 day post-amputation (*p*=0.02) and a significant increase in S/G2-AxFUCCI starting at 3 days post-amputation (*p* = 0.02). One-way ANOVA followed by Tukey’s HSD test revealed a significant increase in Transition-AxFUCCI cells at 1 and 2 days post-amputation (*p*=0.04 and 0.02 respectively), but not at later days post-amputation (*p* > 0.99). Mitotic cells were absent in the 0 day samples, precluding statistical analysis. The percentage of AxFUCCI-negative cells was negligible and did not change at any time point (*p* = 0.40, Wilcoxon rank sum). Each dot represents data from one tail. Error bars indicate standard deviations. ns: not statistically significant. **C) The emergence of Transition-AxFUCCI cells at 1 and 2 days post-amputation was confirmed in tissue sections.** Images are single plane confocal images of 10 μm-thick tissue cross sections fixed at the indicated times after amputation. Percentages indicate the percentage of Transition-AxFUCCI cells at each time point (mean ± standard deviation). *n* = 3 tails per time point, ~ 400 cells counted in each, corresponding to ~ 750 μm of spinal cord. **D) Transition-AxFUCCI-expressing cells at 1 and 2 days post-amputation reside at the G1/S transition.** Transition-AxFUCCI cells can be either at the G1/S transition or the G2/M transition. Following an 8 hour (0 days) or 24 hour (1 and 2 days) EdU pulse, only G2/M cells should become labelled with EdU. Tails were fixed, then processed for EdU detection and tissue sectioning. > 97 % of Transition-AxFUCCI cells were EdU-negative at 1 and 2 days post-amputation, indicating that they reside at the G1/S transition. *n* = 3 tails per time point. A total of 36, 442 and 315 Transition-AxFUCCI cells were assayed at 0, 1 and 2 days post-amputation respectively. **E) Cell cycle dynamics during axolotl spinal cord regeneration**.

After amputation, we detected a significant drop in G0/G1-AxFUCCI-expressing ependymal cells already at day 1. Reciprocally, the number of S/G2-AxFUCCI-expressing cells increased significantly starting at 3 days and reached 50 % at days 4 and 5 post-amputation, compared to a baseline percentage below 10 % (Figures 5A and B). M phase cells started to appear noticeably from day 4, although it was not possible to perform statistical analysis due to the absence of M phase cells in the 0 day post-amputation samples (Figure 5B). Intriguingly, we observed a transient and short ‘burst’ of Transition-AxFUCCI cells at days 1 and 2 post-amputation. The percentage of Transition-AxFUCCI cells increased to 21 ± 12 % and 23 ± 12 % respectively during this burst, before declining back to the baseline level of 5 ± 4 % at 3 days (Figure 5B). We confirmed the accuracy of these quantifications by preparing and quantifying tissue sections from replicate AxFUCCI spinal cords (Figure 5C and Figure 5-figure supplement 1A).

Transition-AxFUCCI could correspond to G1/S transition or G2/M transition (Figure 3A). We hypothesized that Transition-AxFUCCI cells at 1 and 2 days post-amputation resided at the G1/S transition, as they appeared between the decline in G0/G1-AxFUCCI cells at day 1 and the increase in S/G2-AxFUCCI cells at day 3. To confirm this, we subjected AxFUCCI animals to an EdU pulse for 24 hours immediately prior to tail harvesting at 0, 1 or 2 days post-amputation. In this assay, Transition-AxFUCCI cells that incorporate EdU should be in G2/M, while those that do not incorporate EdU should be in G1/S (Figure 5-figure supplement 1B). Consistent with our expectations, Transition-AxFUCCI cells at 1 and 2 days post-amputation were almost entirely EdU-negative and therefore resided in G1/S (Figure 5D and Figure 5-figure supplement 1C,D).

In sum, AxFUCCI revealed the following cell cycle dynamics during regeneration (summarized in Figure 5E). Most ependymal cells in the uninjured spinal cord reside in G0/G1 phase of the cell cycle. Following tail amputation, ependymal cells start to leave G0/G1 within the first day of amputation, transit through G1/S at days 1 and 2, enter S/G2 from day 3 onwards and undergo mitosis from day 4. The fact that these behaviors are readily observed at the population level indicates a high level of synchrony among ependymal cells in the recruitment zone. The G1/S transition acts as a discrete landmark in the cell cycle at which this synchrony can be inferred reliably. Transition-AxFUCCI cells are very rare (~ 5 %) at amputation. We take the 4.5-fold increase in Transition-AxFUCCI-expressing cells at days 1 and 2 post-amputation as a strong indication of cell cycle synchrony during early spinal cord regeneration, a key prediction of our model.

### A convergence in regenerative response from distinct baselines

The AxFUCCI quantifications validated key predictions of our model in terms of the size of the ependymal cell recruitment zone and in demonstrating high cell cycle synchrony during the regenerative response. We were intrigued that these agreements occurred despite a significant difference in starting cell cycle conditions (day 0) between the AxFUCCI animals and our model, which was based on data described in (Rodrigo Albors *et al*. 2015). The AxFUCCI animals were of size 5.5 cm from snout to tail and, on the day of amputation, 11 ± 6 % of ependymal cells expressed S/G2-AxFUCCI. In contrast, our model was parametrized using measurements acquired from 3 cm, non-transgenic axolotls, in which the baseline percentage of S-phase cells was four-fold higher, as inferred from cumulative BrdU labeling (Rodrigo-Albors *et al*., 2015). Despite these differences, we found the speed of spinal cord tissue regeneration to be similar in the two datasets (Figure 3-figure supplement 4B). This could reflect a convergence in regenerative response at the cell cycle level.

We investigated further the kinetics of this convergence by using our model to generate day-by-day simulations of the spatial distribution of cells in G1 or S phase and comparing these simulations to the cell cycle phase quantifications made from the AxFUCCI animals (Figure 6 A-D, Video 5). We validated the simulations by testing where and when ependymal cells would leave G1 phase to enter S phase and found that, as expected, simulated G1 to S transitions were more frequent posterior to the experimentally derived recruitment limit (Figure 6 E), which at day 4 and 5 overlap with the determined AP borders. Consequently, simulated cell divisions (M phase) exhibit a similar pattern (Figure 6 – figure supplement 1).

**Figure 6.**
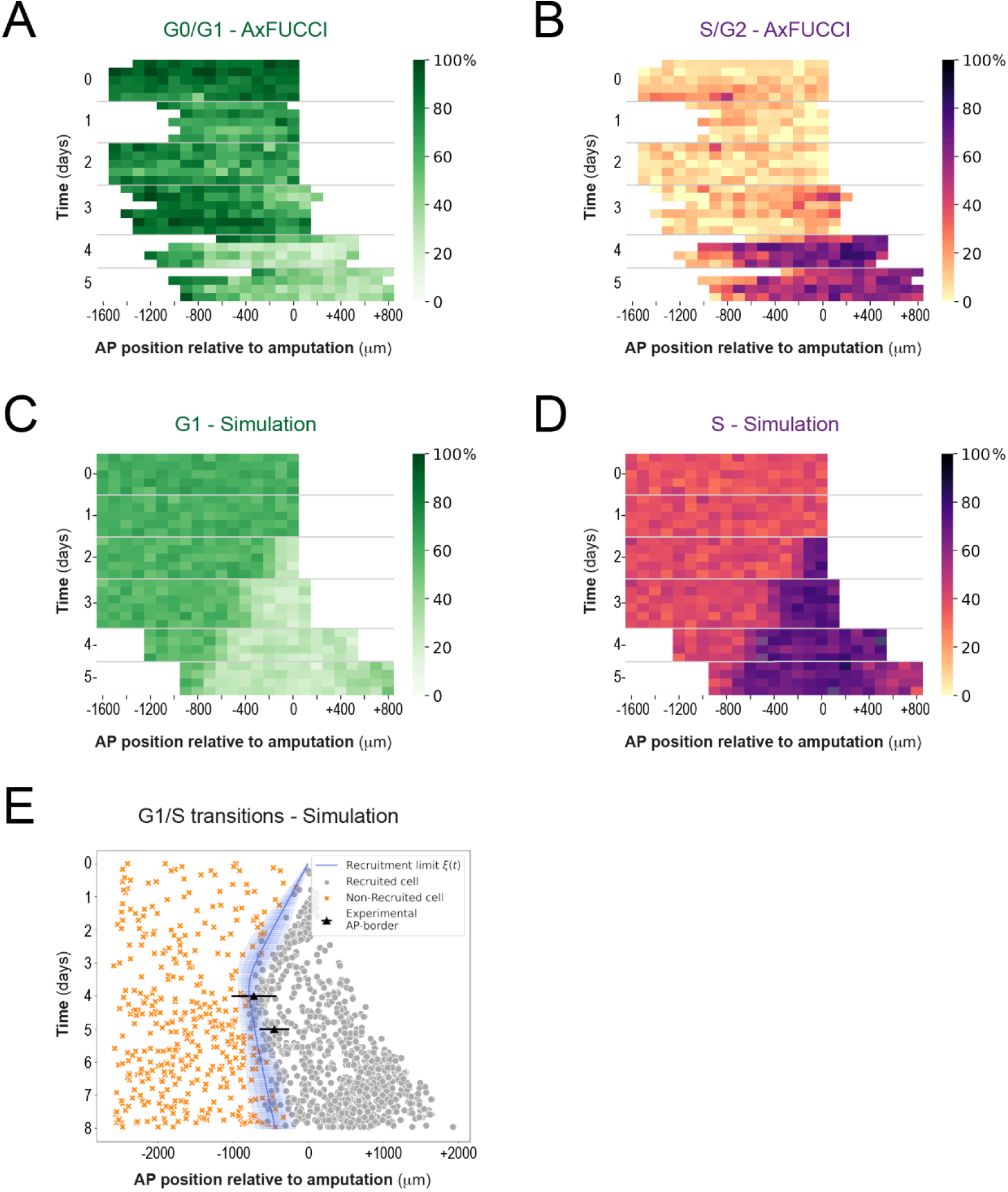
Convergence of cell cycle response between the model simulations and the AxFUCCI data. Heatmaps depicting spatiotemporal distribution of G0/G1-AxFUCCI cells **(A)**, S/G2-AxFUCCI cells **(B)**, model-predicted cells in G1 phase **(C)** and S phase **(D)**. x-axes depict AP position with respect to the amputation plane (AP position = 0 μm). y-axes depict time (days) post-amputation and both experimental and simulated replicates within each timepoint. The color codes correspond to the percentage of cells in the corresponding phase. Transition-AxFUCCI data are in Figure 6 – figure supplement 2. **(E)** Model-predicted occurrence of G1-S transitions is more often within the recruited (grey circles) compared to the non-recruited (orange crosses) cells. At days 4 and 5, the recruitment limit overlap with the AP borders. Independent simulations from 10 independent seeds are shown and the model is parameterized as in Figure 2.

We compared the model simulations to the AxFUCCI data. As noted, the cell cycle conditions at day 0 differ between the AxFUCCI experiments and our model but match quantitatively at day 4 post-amputation (Figure 6 A-D, Video 5). We found the appearance of the recruitment zone to be later in the AxFUCCI animals (days 4-5) than in the simulations (day 2). However, once the recruitment zone is evident in the AxFUCCI data, its size at day 4 and 5 is comparable to the one predicted by the simulations at those times. Thus, the recruitment zone in the AxFUCCI spinal cord manifests more synchronously and rapidly than in the simulations, in which it increases gradually in size in a posterior-to-anterior direction from days 2 to 4. This is likely a consequence of the larger pool of G0/G1 cells in the AxFUCCI animals at days 0-2 compared to the model, which incurs a collective (but relatively synchronous) lag in S-phase entry (recruitment zone manifestation). Our data reveal two contrasting trajectories towards achieving a common regenerative output.

## Discussion

The tissue response to spinal cord injury differs greatly across vertebrates. In mammals, including humans, injuries to the spinal cord often result in permanent tissue damage. In salamanders like the axolotl, however, the ependymal cell response is tightly orchestrated to faithfully rebuild the missing spinal cord (Joven & Simon, 2018; Tazaki, Tanaka & Fei, 2017). Following tail amputation, ependymal cells in the axolotl spinal cord switch from slow, neurogenic to faster, proliferative cell divisions (Rodrigo Albors *et al*., 2015). These faster cell cycles lead to the expansion of the ependymal/neural stem cell pool and drive an explosive regenerative outgrowth. However, the mechanisms regulating cell cycle dynamics during regeneration are not fully understood. Here, by using a modelling approach tightly linked to experimental data, we find that the spatiotemporal pattern of cell proliferation in the regenerating axolotl spinal cord is consistent with a signal that propagates anteriorly 828 um from the injury site during the first 85 hours post-amputation. Although, for simplicity, we refer in this manuscript to a single amputation-induced signal, our model could naturally extend to the combined output of multiple genetic, chemical and/or biophysical signals. Moreover, we show that shortening of S phase is sufficient to explain the explosive growth observed during the first days of regeneration, but that both S and G1 shortening are necessary to explain/sustain further outgrowth before the first new-born neurons are seen (Rodrigo Albors *et al*., 2015).

Compared to the number of mathematical models designed to unveil pattern formation phenomena during development (Morelli *et al*., 2012), modelling in regeneration is still in its infancy (Chara *et al*., 2014). An interesting example of modelling applied to regenerative processes was given by a system of deterministic ordinary differential equations that was superbly used to disentangle how secreted signaling factors could be used to control the output of multistage cell lineages in a self-renewing neural tissue, the mammalian olfactory epithelium (Lander *et al*., 2009). Another mathematical model based on ordinary differential equations was conceived to establish the causal relationship between the individually quantified cellular processes to unravel the stem cell dynamics in the developing spinal cord in chick and mouse (Kicheva *et al*., 2014). In a similar approach, we previously modelled the regenerating axolotl spinal cord by means of a system of deterministic ordinary differential equations describing the kinetics of the cycling and quiescent ependymal cell numbers which we mapped to a model of spinal cord outgrowth (Rost *et al*., 2016). This allowed us to conclude that while cell influx and cell cycle re-entry play a minor role, the acceleration of the cell cycle is the major driver of regenerative spinal cord outgrowth in axolotls (Rost *et al*., 2016). A more recent study based on ordinary and partial differential equations involving cell proliferation was used to predict the spinal cord growth of the knifefish (Ilieş, Sipahi & Zupanc, 2018).

In this study, we investigated the spatiotemporal distribution of cell proliferation during axolotl spinal cord regeneration. To do so, and in contrast with the aforementioned articles, we developed a more general and yet accurate cell-based model introducing the spatial dimension relevant to the problem: the AP axis. To further build a more realistic model, we included non-deterministic attributes: an exponential distribution of the initial coordinates along the cell cycle and a lognormal distribution of the cell cycle length. In the model, a signal shortens the cell cycle of ependymal cells along the AP axis as a consequence of shortening their G1 and S phases, as we reported earlier (Rodrigo Albors *et al*., 2015). Regulation of G1 and S phases are well-known mechanisms controlling cell fate and cell output in a number of developmental contexts. In the brain, G1 lengthening results in longer cell cycles in neural progenitors undergoing neurogenesis (Lukaszewicz *et al*., 2005; Calegari *et al*., 2005; Takahashi, Nowakowski & Caviness, 1995), while experimental shortening of G1 in neural progenitors of the cerebral cortex results in more proliferative divisions, increasing the progenitor pool and delaying neurogenesis (Salomoni & Calegari, 2010; Lange, Huttner & Calegari, 2009; Pilaz *et al*., 2009; Calegari, F. & Huttner, 2003). Here, we have shown that the shortening of G1 during spinal cord regeneration is necessary to sustain the expansion of the ependymal cell pool. Together, these findings point to the regulation of G1 length as a key mechanism regulating the output of neural stem/progenitor cell divisions in development and in regeneration. The length of S phase is also regulated during development by modulating the number of DNA replication origins (Nordman & Orr-Weaver, 2012). In mammals, shortening of S phase seems to play a role in regulating the mode of cell division: mouse neural progenitors committed to neurogenesis and neurogenic cortical progenitors in the ferret undergo shorter S phase than their self-renewing/proliferative counterparts (Turrero García *et al*., 2016; Arai *et al*., 2011). In the axolotl, regenerating ependymal cells shorten S phase during the expansion/outgrowth phase. Together, these findings suggest that the regulation of S phase controls cell output in the context of development and regeneration rather than influence the mode of cell division. The combined shortening of S and G1 in the regenerating spinal cord sustains the expansion of the resident ependymal/neural stem cell pool at the expense of neurogenesis. In this line, experimentally shortening G1 and S phases in cortical progenitors of the developing mouse brain delayed the onset of neurogenesis (Hasenpush-Theil *et al*., 2018). Our findings add to the evidence that cell cycle regulation is a key mechanism controlling the number and type of cells needed to generate and regenerate a tissue.

Another prediction of our model is that a signal must spread about 800 μm from the injury site while recruiting ependymal cells 85 hours after amputation to explain the spatiotemporal pattern of cell proliferation in the regenerating spinal cord. In order to test this prediction experimentally, we adapted FUCCI technology to axolotls, which enabled us to visualize cell cycle dynamics *in vivo*. We found remarkable agreement between our prediction and the size and timing of appearance of the recruitment zone in AxFUCCI spinal cords. Our prediction was made based on data from 3 cm snout-to-tail axolotls, while the AxFUCCI measurements were taken from 5.5 cm axolotls. That the size of the recruitment zone is constant between these two animal sizes could be important in understanding the identity of the injury-induced signal and how it spreads to recruit ependymal cells. Future experiments will determine if the size of the recruitment zone also remains constant in even larger axolotls.

A characteristic feature of our model is that G1 shortening after amputation causes ependymal cells to partially synchronize with one another as they pass through G1. Cell cycle synchronization is difficult to measure in cells *in vivo*. Here, AxFUCCI’s property of labelling short, discrete landmarks in the cell cycle (*e.g*. G1/S transition) enabled us to visualize high G1/S synchrony at 1 and 2 days post-amputation *in vivo*. It will be interesting to assess whether a similar phenomenon occurs during regeneration of other tissues in the axolotl (*e.g*. limb) and in other regenerative organisms.

Although we observed an excellent match between our model simulations and the AxFUCCI data in terms of recruitment zone size at days 4 and 5 post-amputation, we also encountered quantitative differences at earlier time points. In particular, we found that significantly fewer ependymal cells were in S/G2 at baseline conditions in the present study compared to our previous study (Rost *et al*., 2016). The AxFUCCI experiments reported here were carried out under COVID-19-related operational restrictions – in particular, animal feeding frequency was reduced. A dietary reduction could plausibly impact baseline ependymal cell proliferation rate and animal growth. We also cannot exclude an impact from general housing conditions, as the previous experiments were performed in a different animal facility with, for example, a different water supply. From the data analysis side, it is important to note that in the AxFUCCI experiments, G0/G1 AxFUCCI cells become Transition AxFUCCI cells and then S/G2 AxFUCCI cells. This means that transition cells could be either cells in the late G1 or in the early S phases. Modeled cells, in contrast, go straight from G1 to S phase. Consequently, we cannot quantitatively equate the ‘Transition phase’ cells between experiments and model. This is why the AxFUCCI data and the model simulations can only be qualitatively compared, especially at days 1 and 2, when transition cells peak. Despite these considerations, we found that the proportions of ependymal cells in the G0/G1 *vs* S/G2 AxFUCCI data at 4 and 5 days post-amputation – *i.e*. during the first regenerative cell cycle – accurately and quantitatively matched the simulations of our model. Moreover, the outgrowth rate of the regenerating spinal cord was consistent between the AxFUCCI animals in this study and the animals measured in our previous study (Figure 3-figure supplement 4B). This is to say that axolotls mount a remarkably consistent regenerative response within their first cell cycle after amputation, possibly converging their cell cycle responses despite differences in baseline. It will be fascinating to investigate the molecular mechanisms that enable this consistent regenerative response in axolotls across age/size and nutrition availability.

An important question now is whether the spatiotemporal cell cycle response observed in this study agrees with known signaling events operating during spinal cord regeneration. A strong candidate molecule for recruiting ependymal cells is the axolotl MARCKS-like protein (AxMLP), a secreted factor involved in the proliferative response during axolotl appendage regeneration (Sugiura *et al*., 2016). AxMLP is normally expressed in spinal cord cells but is upregulated following tail amputation, peaking 12 to 24 h after amputation and returning to basal levels a day later (Sugiura *et al*., 2016). The timing prediction of our model is in agreement with the peak of AxMLP followed by a downstream period of signal decoding to instruct intrinsic cellular changes that lead to faster cell cycles. Moreover, the secreted nature of AxMLP protein could explain the long-range proliferative response in the regenerating spinal cord. In the future, a tighter time-course characterization of AxMLP localization throughout axolotl spinal cord regeneration will help to put our predictions to test.

Changes in the biophysical properties of the amputated tail could also trigger the orderly increase in cell proliferation. In Xenopus tadpoles, tail amputation leads to the activation of the H+ V-ATPase which is necessary and sufficient to promote tail regeneration (Adams *et al*., 2007). In the axolotl, tail amputation triggers changes in calcium, sodium, and membrane potential at the injury site (Ozkucur *et al*., 2010) while spinal cord transection induces a rapid and dynamic change in the resting membrane potential which drives a c-Fos dependent gene expression program promoting a pro-regenerative response (Sabin *et al*., 2015). The proliferation-inducing signal could also be of mechanical nature (Chiou & Collins, 2018). In this direction, it is interesting that spinal cord transection in the zebrafish induces an immediate alteration in the mechanical properties in the lesion site, which gradually returns to normal (Schlüßler *et al*., 2018). Our predictions of the temporal and spatial distribution that such proliferation-inducing signal could have will guide efforts to narrow down the mechanisms responsible for successful spinal cord regeneration.

Taken together, our study provides a finer mechanistic understanding of the cell cycle kinetics that drive spinal cord regeneration in axolotl and paves the way to search for the signal or signals that launch the successful ependymal cell response to spinal cord injury.

## Materials and Methods

### 1. Computational methods

#### 1.1 Model of developing and regenerating axolotl spinal cord

We modeled the spinal cord as a densely packed row of ependymal cells. Since all the cells are assumed identical rigid spheres, the model effectively involves only one spatial dimension: the anterior-posterior (AP) axis of the spinal cord. We assumed that cells are either cycling or quiescent, where the fraction of cycling cells is the growth fraction *GF*. We considered that each cycling cell *i* located in the position *x_i_* at the time *t* proliferates with a certain random cell cycle length lognormally distributed *T_i_* (*x_i_, t*) and has a certain age within its cell cycle, *C_i_* (*x_i_, t*), defined as a coordinate along the cell cycle or clock (0 ≤ *C_i_* (*x_i_, t*) < *T_i_* (*x_i_, t*)). In the initial condition, each cycling cell has a random age 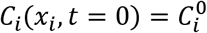 along its particular cell cycle length *T_i_* (*x_i_, t*), where the 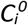 distribution is given by 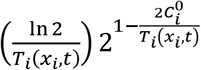. As time *t* goes by, each cycling cell increases its clock *C_i_* (*x_i_, t*) deterministically until it reaches its corresponding *T_i_* (*x_i_, t*) value. At this precise moment, the cell divides and one daughter inherits its mother’s AP coordinate while the other is intercalated between the first daughter and the posterior neighboring cell. This last feature of the model is the implementation of what we earlier defined as “cell pushing mechanism” (Rost *et al*., 2016). After cell division, the daughter cells reinitiate their clocks and *c_i_*(*x_i_, t*) = 0. This model predicts that after a time of approximately one cell cycle length, mitotic events will occur along the AP axis, contributing to the growth of the spinal cord during development (Figure 1A).

To study the evolution of the tissue under a regenerative setup, we focused on the tissue response to an amputation modeled by simply removing the most posterior cells. We modeled the regenerative response in the remaining *N_0_* cells by assuming that amputation triggers the release of a signal, which spreads with constant velocity anteriorly over the AP axis while recruiting the ependymal cells. We assumed that cell recruitment stops at time *τ*, rendering *λ* μm of cells anterior to the amputation plane recruited and a recruitment velocity-*λ*/*τ* during the interval 0 ≤ *t* ≤ *τ*. We notated the AP position of the most anterior cell recruited by the signal as *ξ*(*t*), the recruitment limit, such that *ξ*(*t* = *τ*) = −*λ*.

Because regenerative ependymal cells shorten G1 and S phases without altering G2 and M phases leading to an acceleration of the cell cycle (Rodrigo Albors *et al*., 2015), we assumed that the signal-induced recruitment instructs regenerating ependymal cells precisely to reduce G1 and S phases, effectively shortening their cell cycle (Figure 1C). We represent here as *G1^long^* and *S^long^ (G1^short^* and *S^short^)* the length of the corresponding phases for ependymal cells of uninjured animals (regenerating animals). We notate with *T^long^* and *T^short^* to the cell cycle length of the ependymal cells in uninjured and regenerating axolotls, respectively.

Note that a cycling cell *i* whose position *x_i_* is anterior to the recruitment limit (*x_i_* < *ξ*(*t*)) is not recruited at time *t* and has a cell cycle length *T_i_* (*x_i_,t*) equal to *T^long^*, that is, continue cycling slowly during the simulations (Figure 1-figure supplement 1B). In contrast, a cycling cell *i* whose position *x_i_* is posterior to the recruitment limit (*x_i_* ≥ *ξ*(*t*)) within the time interval 0 < *t* ≤ *τ* is irreversibly recruited and consequently has a cell cycle length *T_i_* (*x_i_,t*) equal to *T^short^*. The progeny of the recruited cells (non recruited cells) have a cell cycle length extracted from the same lognormal distribution of *T^short^* (*T^long^*) (Figure 1C).

We assumed that recruitment of a cell *i* located in the position *x_i_* at time *t* induces an irreversible transformation in its cell cycle coordinate *C_i_* (*x_i_, t*) → *C_i_*’ (*x_i_, t*), where *C_i_* (*x_i_, t*) and *C_i_*’ (*x_i_, t*) are the original and transformed cell cycle coordinates, respectively. This means that the cell cycle coordinates of these cells undergo an irreversible coordinate transformation, modifying their cycling according to the cell cycle phase in which they are in at the moment of recruitment, as we describe the following subsections (Figure 1-figure supplement 1B):

##### 1.1.1 When the cells are in the G1 phase at the time of recruitment

We assumed that if at the moment of amputation *t*, a cell *i* would be in a cell cycle coordinate *C_i_* (*x_i_, t*) within 0 ≤ *C_i_*(*x_i_, t*) ≤ *G*1^*long*^ — *G*1^*short*^, the new cell cycle coordinate is as follows (Figure 1C and Figure 1 – figure supplement 2):

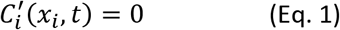

which would induce a synchronization. In contrast, if at the moment of amputation *t*, a cell *i* would be in a cell cycle coordinate *C_i_* (*x_i_, t*) within *G*1^*long*^ — *G*1^*short*^ ≤ *C_i_* (*x_i_ t*) ≤ *G*1^*long*^, the new cell cycle coordinate is:

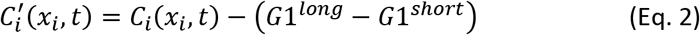

That is, these cells continue cycling as before. Taken together, the cells in G1 result partially synchronized (Figure 1C and Figure 1 – figure supplement 2).

##### 1.1.2 When the cells are in the S phase at the time of recruitment

Taking into account that in S phase all DNA must be duplicated for cell division to occur, we considered a different mechanism to model S phase shortening based on proportional mapping. The new cell cycle coordinate of this cell are proportionally mapped to the corresponding coordinate of a shortened S phase in the next simulation step. Thus, we assumed that if at the moment of amputation *t*, a cell *i* would be in the S phase, that is, in a cell cycle coordinate *C_i_*(*x_i_, t*) within *G*1^*long*^ ≤ *C_i_*(*x_i_, t*) ≤ *G*1^*long*^ + *S^long^*, the transformed cell cycle coordinate relative to the S phase length is invariant (Figure 1C and Figure 1 – figure supplement 2):

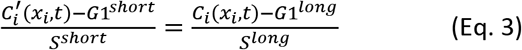

As a consequence, the transformed cell cycle coordinate is as follows:

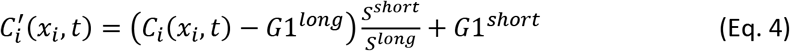

##### 1.1.3 When the cells are either in the G2 or M phase at the time of recruitment

We previously demonstrated that sum of G2 and M phase lengths of the ependymal cells of axolotl spinal cords were conserved after amputation (Rodrigo Albors *et al*., 2015). Hence, once a cell *i* is in the joint G2 + M phases, the remaining time to complete the cell cycle is the same for both, the original and the transformed cell cycle coordinates (Figure 1C):

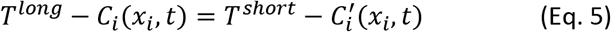

As a consequence, the transformed cell cycle coordinate can be calculated as follows:

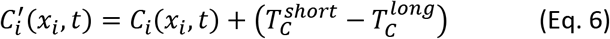

##### 1.1.4 When the cells are in the G0 phase at the time of recruitment

We assumed that if a recruited cell *i* is quiescent at the moment *t* of recruitment, that is, in G0, it progress from this phase to the short-G1 phase after a certain delay *t_G0-G1_*.

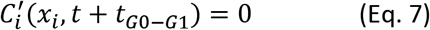

#### 1.2 Model parametrization

The model parameters are summarized in Table 1. Briefly, the ependymal cell length along the AP axis, the distributions of cell phases durations and growth fraction were fixed from our previous publication (Rodrigo Albors *et al*., 2015). The only free model parameters are the remaining anterior cells after amputation *N_0_*, the maximal length *λ* along the AP axis of the putative signal and *τ*, the maximal time of cell recruitment.

#### 1.3 Fitting procedure of the experimental switchpoint with the theoretical recruitment limit *ξ*(*t*)

The experimentally obtained switchpoint of the regenerating axolotl spinal cord (extracted from Rost *et al*., 2016) was fitted with the model-predicted recruitment limit *ξ*(*t*). We followed an Approximate Bayesian Computation method (ABC) to estimate the distribution of the parameters that better reproduces the experimental switchpoint data by our recruitment limit model. The ABC methods bypass the requirement for evaluating likelihood functions and capture the uncertainty in estimates of model parameters (Csilléry *et al*., 2010). In particular, we used pyABC (Klinger, Rickert & Hasenauer, 2018), a high-performance framework implementing a Sequential Monte Carlo scheme (ABC-SMC), which provides a particularly efficient technique for Bayesian posterior estimation (Toni *et al*., 2018).

Briefly, we generated a series of stochastic simulations from the model (described in the previous section) from sampled points of the parameter space. Each fitting run was initialized with a population size of 1000 samples. All the prior distributions of the parameters were defined as a discrete uniform distribution: *N_0_* ~ unif{100, 300}, *λ* ~ unif{500, 1500} and *τ*~ unif{1,192}. Where the limits of *τ* were basically given by the entire experimental observation time (8 days) in hours. The limits of *λ* (in μm) and *N_0_* were initially estimated by previous simulation trials.

The sampled parameter values were accepted only when the distance function *d* between simulated recruitment limit and experimental switchpoint was lower than a given tolerance *ε*. The distance function was defined as follows:

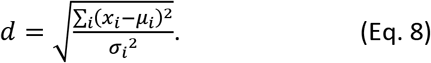

where *μ_i_*, *σ_i_*, and *x_i_*, correspond to the mean, standard deviation of the experimental switchpoint and the simulated recruitment limit, respectively, at the experimental time points *i* (4, 6 and 8 days). At each iteration, the parameter distributions were updated and re-sampled. The new tolerance *ε* was then calculated as the median of the distances from the last accepted sample population. The outcome of the algorithm was a sample of parameter values inferring their posteriors distributions (Figure 2–figure supplement 1A,B). Convergence of the method was assessed by following the value of *ε* and the acceptance rate, defined as the accepted number of simulations divided by the total number of simulations at each iteration step (Figure 2–figure supplement 1C,D).

#### 1.4 Clones trajectories and velocities

We calculated the clone trajectories following the positions of each clone in random simulations. When a cell divided, we kept the mean position of the clone cells as the clone position. In Figure 2-figure supplement 2, a total of 11 tracks are shown, the first trajectory starts at 0 (the amputation plane) and the last at −1100 μm (with a sampling of 50 μm, approximately). To estimate the mean velocity of clones at different spatial positions in this model, the space along the AP axis was subdivided into 800 μm bins. For each clone trajectory, the positions were grouped according to these bins. Groups containing less than two measurements were excluded. The average clone velocity for each group was estimated with linear regression. Then, the mean and standard deviation of the velocity of all the clones in a bin was calculated.

#### 1.5 Coordinate system

In all our simulations, the time starts with the event of amputation. Space corresponds to the anterior-posterior (AP) axis, where 0 represents the amputation plane and positive (negative) values are posterior (anterior) locations.

#### 1.6 Model implementation and computational tools

The models were implemented in Python 3.0. Simulations and data analysis were performed using Numpy (Oliphant, 2006) and Pandas (McKinney, 2010) while data visualization was executed with Matplotlib (Hunter, 2007).

#### 1.7 Supplementary notebooks

Jupyter Notebook (http://jupyter.org/) containing the source code for all computations performed and referred to as Cura Costa *et al*., 2021 in this study can be found at https://doi.org/10.5281/zenodo.4557840.

### 2. Experimental materials and methods

#### 2.1 Molecular biology

AxFUCCI plasmid was constructed by standard restriction cloning. Relevant features of the AxFUCCI plasmid are: (i) SceI site for stable transgenesis; (ii) CAGGs synthetic promoter for ubiquitous expression; (iii) G0/G1-AxFUCCI probe (axolotl Cdt1[aa1-128]-GSAGSAAGSGEF glycine/serine linker-mVenus); (iv) T2A ‘self-cleaving’ viral peptide; (v) S/G2-AxFUCCI probe (mCherry-SGGGGGSGGGGS glycine/serine linker-axolotl Gmnn[aa1-93]); (vi) rabbit beta-globin polyadenylation sequence; (vii) SceI site for stable transgenesis. PCR products were amplified using the primers listed in Table 2 and ligated in two rounds into a vector already harboring SceI sites, a CAGGs promoter, a T2A sequence followed by mCherry and rabbit beta-globlin polyadenylation sequence.

**Table 2.**
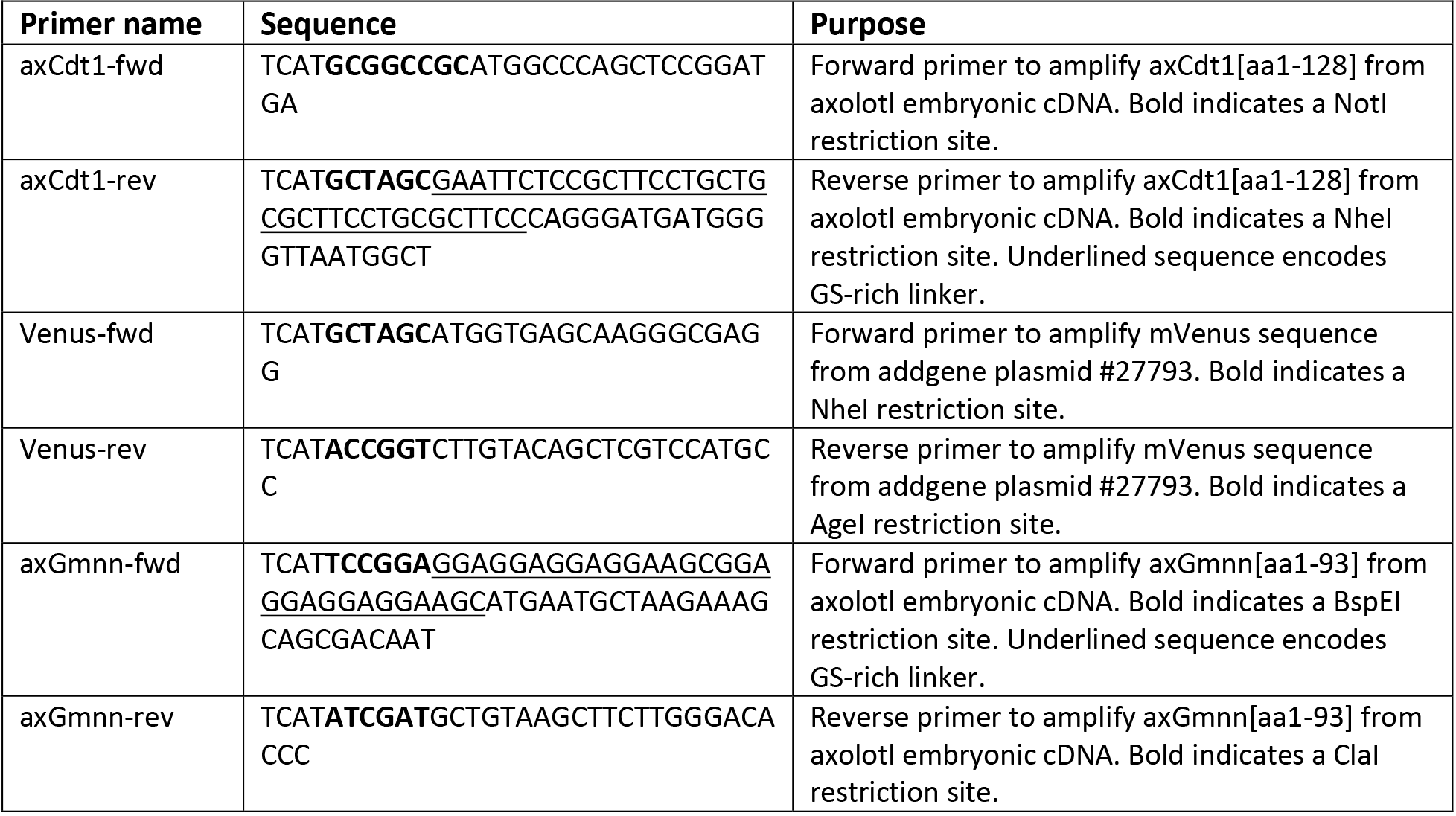
Primers used in this study.

Primers were purchased as 20μM stocks (Sigma-Aldrich, standard de-salt). AxFUCCI plasmid sequence was verified by Sanger sequencing.

#### 2.2 AL1 cell culture

The immortalized axolotl ‘AL1’ cell line was grown in a humidified incubator at 25°C, with 2% CO_2_. The cell culture medium contains: 62.5% DMEM, 10% Fetal Bovine Serum, 25% water, supplemented with 100U Penicillin-Streptomycin, Glutamine, Insulin. AL1 cells were passaged every week at a ratio of 1:2 into gelatin-coated flasks.

#### 2.3 AL1 cell electroporation and live imaging

Electroporation was performed using a Neon Transfection System (Thermo Fisher Scientific). 50,000 AL1 cells were electroporated with 1μg of AxFUCCI plasmid in 70% PBS/water using the following settings: 750V, 35ms pulse width, 3 pulses. Electroporated AL1 cells were plated onto a glass-bottomed Ibidi imaging dish coated with gelatin. After two days, the cell culture medium was exchanged, and the dish placed in a Celldiscoverer 7 automated live cell imaging microscope chamber (Zeiss). The microscope chamber was maintained at 25°C, with 2% CO_2_. Cells were imaged hourly over the course of 7 days for Venus and mCherry fluorescence, and brightfield.

#### 2.4 Fluorescence intensity track measurements

AxFUCCI fluorescence intensities were measured using the TrackMate plugin for FIJI (Tinevez, Perry & Schindelin *et al*., 2017). Fluorescence intensities were normalized to the maximum intensity observed for the respective fluorophore during the experiment.

#### 2.5 DNA quantification by flow cytometry

AxFUCCI-electroporated AL1 cells were incubated for 90 minutes with cell culture medium containing 10μg/ml Hoechst DNA stain. After incubation, AL1 cells were washed once with 70% PBS/water, then dissociated into single cells using Trypsin. Dissociation was terminated by adding a 1:1 volume of serum-containing cell culture medium. Cells were pelleted and re-suspended in 70% PBS/water, then filtered through a 50μm cell filter. Cells were analysed for DNA content using a BD LSRFortessa Flow Cytometer and FlowJo software.

#### 2.6 Axolotl husbandry and transgenesis

d/d and AxFUCCI axolotls (*Ambystoma mexicanum*), snout-to-tail length 5.5cm, were raised in individual aquaria. Axolotl breedings were performed by the IMP animal facility. All experiments were performed in accordance with locally applicable ethics committee guidelines and within a framework agreed with the Magistrate of Vienna (Genetically Modified Organism Office and MA58, City of Vienna, Austria). Axolotls were anaesthetized with benzocaine (Sigma) diluted in tap water prior to amputation and/or imaging.

AxFUCCI axolotls were generated by I-SceI meganuclease-mediated transgenesis using previously described methods (Sobkow *et al*., 2006). Briefly, one-cell stage fertilised d/d axolotl eggs were dejellied and injected with 5nl of injection mix (~0.5ng AxFUCCI plasmid and 0.005U I-SceI meganuclease (NEB) diluted in 1X CutSmart buffer (NEB)). Injected axolotl eggs were maintained in 0.1X MMR/tap water at room temperature until screening. Transgenic founder (F0) animals were identified by their Venus fluorescence using an AXIOzoom V16 widefield microscope (Zeiss). F1 germline-transmitted AxFUCCI animals were used for all experiments in this study. Sample sizes were determined empirically and within the confines of experimentation under COVID19 pandemic-induced operational restrictions.

#### 2.7 Tissue clearing and lightsheet imaging

AxFUCCI tails were fixed overnight at 4°C in 4% paraformaldehyde. Fixed tails were washed well with PBS, then de-lipidated for 30 minutes at 37°C in Solution-1 of the DEEP-clear tissue clearing protocol (10% v/v THEED, 5% v/v Triton X-100, 25% w/v urea in water) (Pende & Vadiwala *et al*., 2020). De-lipidated tails were washed with PBS, then incubated for 2 hours in PBS containing 10μg/ml DAPI. Tails were washed well, then incubated overnight in Easyindex refractive index matching solution (LifeCanvas technologies). Samples were kept dark at all times to prevent bleaching of AxFUCCI fluorescence. Cleared AxFUCCI tails were imaged in EasyIndex solution using a LightSheet.Z1 microscope (Zeiss) and custom chamber.

#### 2.8 Preparation of tissue sections

AxFUCCI tails were fixed overnight at 4°C in 4% paraformaldehyde. Fixed tails were washed well with PBS, then incubated overnight in 30% sucrose in PBS. The following day, samples were embedded in OCT (Optimal Cutting Temperature) compound, frozen on dry ice and stored at −80°C until sectioning. Cryosections of 10μm thickness were prepared from frozen blocks and stored at −20°C until use.

#### 2.9 Immunostaining and imaging of tissue sections

Cryosections were warmed up to room temperature, then washed extensively with PBS to remove OCT. Sections were blocked for 2 hours at room temperature with 10% normal goat serum (NGS) diluted in PBS containing 0.2% Triton X-100 (PBTx). Blocked samples were incubated with primary antibody diluted in 1% NGS overnight at 4°C. The following day, sections were washed well with PBTx, then stained with Alexa Fluor-conjugated secondary antibodies diluted 1:500 in PBTx for 2 hours at room temperature. DAPI was included in the secondary staining solution at a concentration of 10μg/ml. Sections were washed well with PBTx and mounted in Mowiol containing 2.5% DABCO (Sigma). The following primary antibodies were used in this study: anti-NeuN (Millipore MAB377, mouse, 1:500), anti-Sox2 (rabbit, 1:1,000, Fei *et al*., 2016). Images were acquired using a LSM980 AxioObserver inverted confocal microscope (Zeiss).

#### 2.10 EdU administration and detection

Anaesthetized axolotls were injected intraperitoneally with 400μM EdU (diluted in PBS) at a dosage of 20μl/g. FastGreen dye (Sigma-Aldrich) was added to the injection mix to aid visualization. Injected axolotls were kept out of water for a 20-minute recovery period under benzocaine-soaked towels. After recovery, injected axolotls were returned to water. Following the desired pulse-chase period, axolotls were sacrificed, tail tissue fixed and cryosections prepared.

EdU detection was performed using the Click-iT 647 EdU detection kit (Thermo Fisher Scientific) according to the manufacturer’s instructions.

#### 2.11 Image analysis

Lightsheet data (AxFUCCI axolotl tails) were digitally re-sectioned or rendered in 3D using Imaris software (Oxford Instruments). For quantification of wholemount data, spinal cords were digitally re-sectioned longitudinally, to yield a continuous strip of spinal cord lumen, and images were exported as TIFFs for cell counting in FIJI. Ependymal cells cell cycle phases were quantified from 25μm thick digital sections. Ependymal cells were defined as cells in direct contact with the spinal cord lumen. Celldiscoverer 7 videos (AL1 cells) were cropped using ZEN blue software (Zeiss), then analyzed using the TrackMate plugin for FIJI, as described above. Tissue section images were analyzed and quantified using FIJI software (Schindelin *et al*., 2012).

#### 2.12 Determining the AP border between two adjacent spatial zones within axolotl regenerating spinal cords

We tested whether ependymal cells in the different cell cycle phases would be heterogeneously distributed along the AP axis of the regenerating spinal cord. To that aim, we fitted the experimental spatial AP profiles of the percentages of G0/G1-AxFUCCI and S/G2-AxFUCCI expressing cells, per animal, with a mathematical model assuming two adjacent homogeneous spatial zones separated by an anterior-posterior border, as follows:

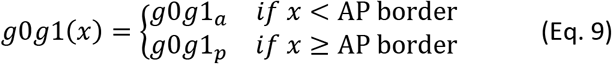

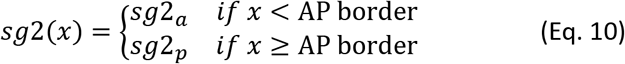

Where *g*0*g*1(*x*) and *sg*2(*x*) are the model variables describing the spatial distribution of G0/G1 and S/G2 cells along *x*, the spatial position along the AP-axis. The model parameters are *g*0*g*1_*a*_, the anterior percentage of G0/G1-AxFUCCI expressing cells, *g*0*g*1_*p*_, the posterior percentage of G0/G1-AxFUCCI expressing cells, *sg*2_*a*_, the anterior percentage of S/G2-AxFUCCI expressing cells, *sg*2_*p*_, the posterior percentage of S/G2-AxFUCCI expressing cells and *AP border*, the border between the anterior and the posterior zones, assumed equal for G0/G1-AxFUCCI and S/G2-AxFUCCI cells.

We fitted the model simultaneously to the AP profile of the percentage of G0/G1-AxFUCCI and S/G2-AxFUCCI expressing cells of each animal and each time by using an Approximate Bayesian Computation method (ABC, see Computational methods section 1.3 for more details of the computational implementation). Each fitting was initialized with a constant population size of 1000 samples. The parameters priors were defined as a discrete uniform between 0 and 100% for *g*0*g*1_*a*_, *g*0*g*1_*p*_, *sg*2_*a*_ and *sg*2_*p*_. The prior for the *AP border* was also a discrete uniform covering all the measured positions along the AP axis. The distance function between the experimental FUCCI data and the two-zones model was defined as:

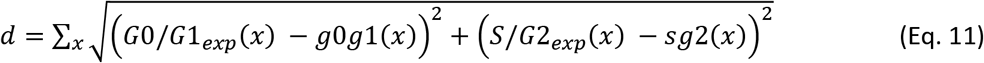

Where *G0/G1_exp_*(*x*) and *S/G2_exp_*(*x*) are the percentage of G0/G1-AxFUCCI and S/G2-AxFUCCI expressing cells, respectively, determined at the bin *x* of the spinal cord AP axis (for each animal and each time).

Each fitting procedure had a total of 30 iterations. Fitting results are shown in Figure 4B, Figure 4-Figure supplement 1 and Figure 4-Figure supplement 2.

For each time, we compared the anterior *versus* the posterior zones of G0/G1-AxFUCCI and S/G2-AxFUCCI data simultaneously by performing a Kolmogorov-Smirnov test between the best-fitting parameters *g*0*g*1*_a_*, *g*0*g*1*_p_ versus sg*2_*a*_, *sg*2_*p*_. Although anterior and posterior zones were indistinguishable from zero to 3 days post amputation, we found a significant difference between anterior and posterior zones in the G0/G1-AxFUCCI and S/G2-AxFUCCI data at day 4 and 5 (Figure 4 – figure supplement 1). The best fitting AP border detected for each time post-amputation are in Figure 4B as vertical gray areas at day 4 and 5, post amputation.

#### 2.13 Statistical analysis and data representation

In Figure 4, Kolmogorov-Smirnov test was implemented by using the Scipy (Virtanen *et al*., 2020) library. Numpy’s (Harris *et al*., 2020) high-level mathematical functions were used all along the simulations and data analysis. In Figure 2, Figure 2 – figure supplement 2, Figure 2 – figure supplement 3, Figure 2 – figure supplement 4 and Figure 3 – figure supplement 4 B were made using Matplotlib (Hunter, 2007) while Figure 4B, Figure 4 – figure supplement 1, Figure 4 – figure supplement 2, Figure 6, Figure 6 – figure supplement 1 and Figure 6 – figure supplement 2 were performed with Seaborn (Waskom *et al*., 2017). In Figure 5 and Figure 5-figure supplement 1, Figure 3-figure supplement 4, statistical analyses were performed using R. AxFUCCI data were tested for assumptions of normality (Shapiro-Wilk test) and equality of variance (Levene’s test) in order to determine the appropriate statistical tests to perform. No data were excluded. Details of statistical tests and their outcomes can be found in the relevant figure legends. Statistical significance was defined as *p* < 0.05. These graphs were plotted using Prism (GraphPad). All the figures were compiled in Adobe Illustrator.

## Acknowledgements

We thank Fabian Rost for critical comments on the manuscript. We also thank the members of the Chara lab and especially Alberto Ceccarelli for interesting discussions. We are grateful to Pietro Tardivo (tissue clearing and imaging), Anastasia Polikarpova (flow cytometry), Alberto Moreno-Cencerrado and Pawel Pasierbek (BioOptics facility, Vienna Biocenter) and the animal caretaker team (Vienna Biocenter), whose support enabled experiments to be performed during the COVID-19 pandemic.

## Supplementary Figures

**Figure 1-figure supplement 1.**
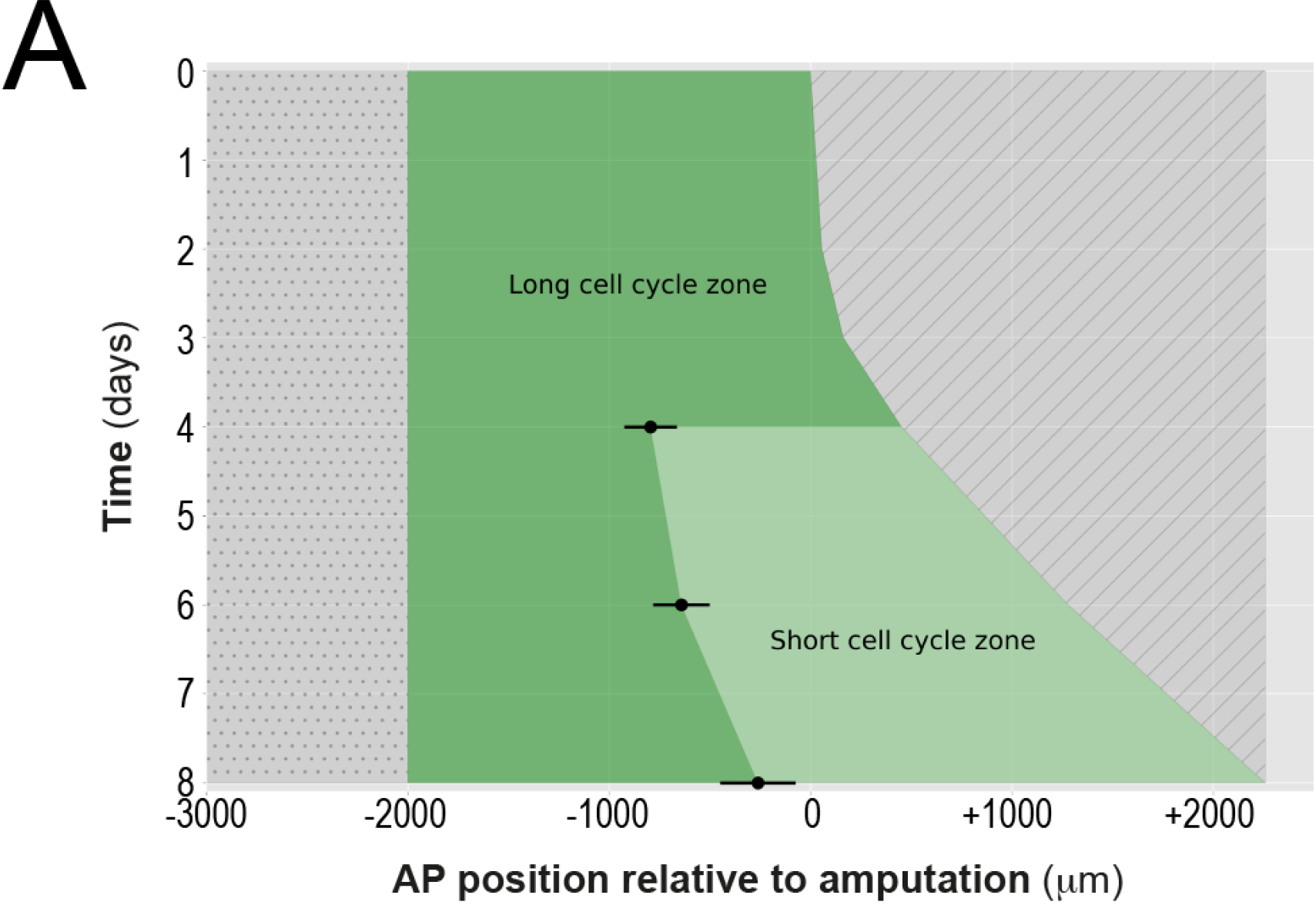
Space-time distribution of cell proliferation during axolotl spinal cord regeneration. Experimental switchpoint (black dots) separating areas of low cell proliferation (Long cell cycle zone, dark-green) from high cell proliferation (short cell cycle zone, light-green) along the anterior-posterior (AP) axis (depicted horizontally) of the axolotl spinal cord. The dashed region marks the space outside of the embryo while the dotted region marks the unaffected part of the embryo (adapted from Fig.2 F’’ of Rost *et al*., 2016). Each switchpoint value was determined as the border separating two spatially homogeneous zones assumed by a mathematical model, which was fitted to the experimental spatial distribution of growth fraction and mitotic index along the axolotl spinal cords (Rost *et al*., 2016).

**Figure 1-figure supplement 2.**
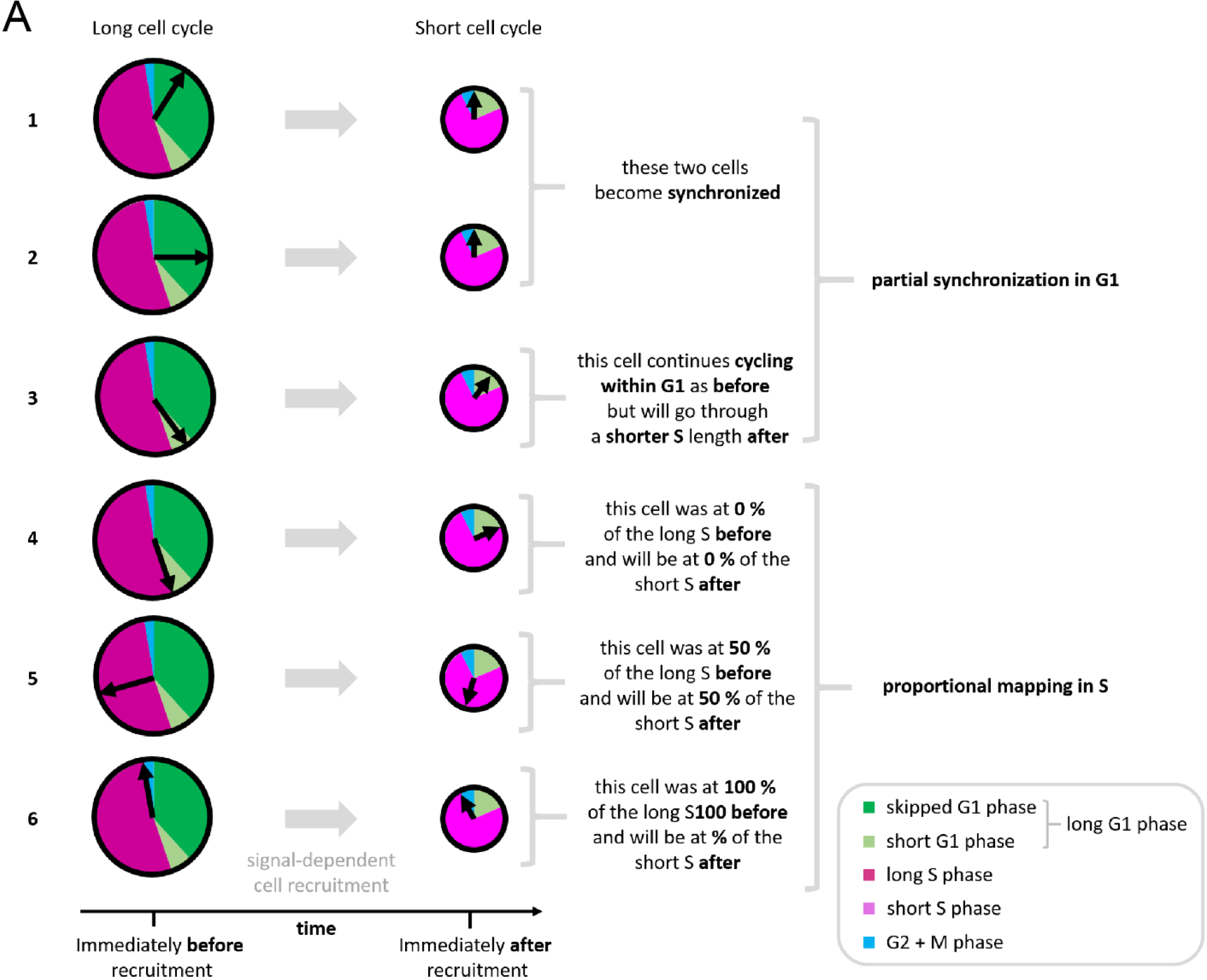
Further explanation of the cell cycle coordinate transformation invoked by the model. Cells in G1 phase at the moment of recruitment can skip the remainder of the phase, depending on where they are located in the long G1 phase. If their cell cycle coordinate is before the difference between the long and the short G1 phases, they skip this initial part of the long G1 phase and synchronize at the beginning of the short G1 phase **(1, 2)**. If their cell cycle coordinate is equal or after the difference between the long and the short G1 phases, they continue cycling as before, but they will go through a short S phase afterwards **(3)**. This generates a partial synchronization mechanism (See computational methods section 1.1.1). If the cells are in S phase at the moment of recruitment, cells follow a proportional mapping **(4 - 4)**. Essentially the cell cycle coordinates within S are scaled into the short S phase (See computational methods section 1.1.2).

**Figure 2-figure supplement 1.**
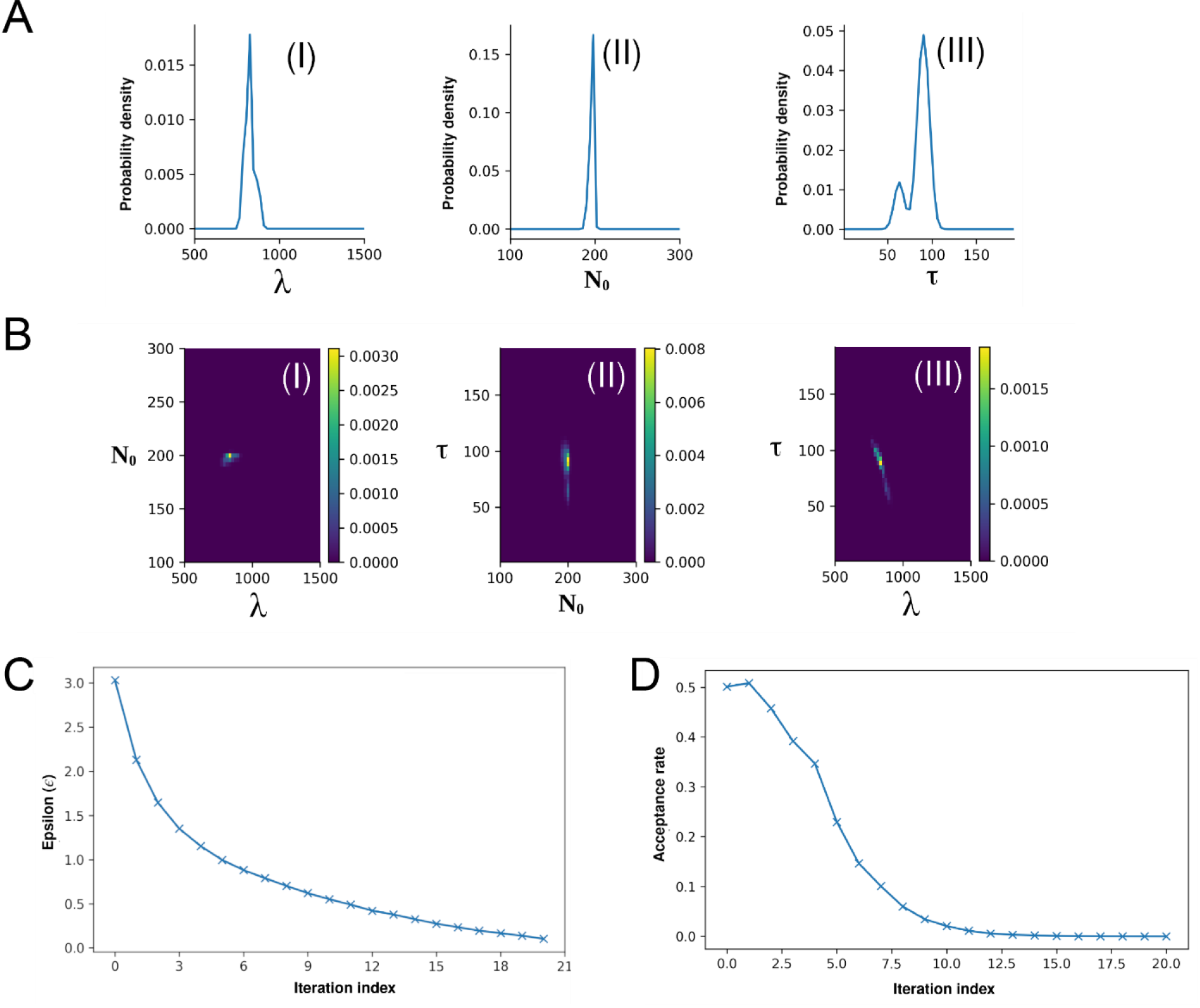
Approximate Bayesian Computation (ABC) fitting of the recruitment limit model to the experimental switchpoint data. **(A)** Posterior distributions of the parameters *λ* (I), *N_0_* (II) and *τ*(III). **(B).** Combined posterior distributions of *N_0_* and *λ* (I), *τ* and *N_0_* (II) as well as *τ* and *λ* (III). **(C and D)** Fitting process convergence depicted by the fitting tolerance *ε* (B) and the acceptance rate (D) *versus* the iteration index. (See computational methods section 1.3 for details).

**Figure 2-figure supplement 2.**
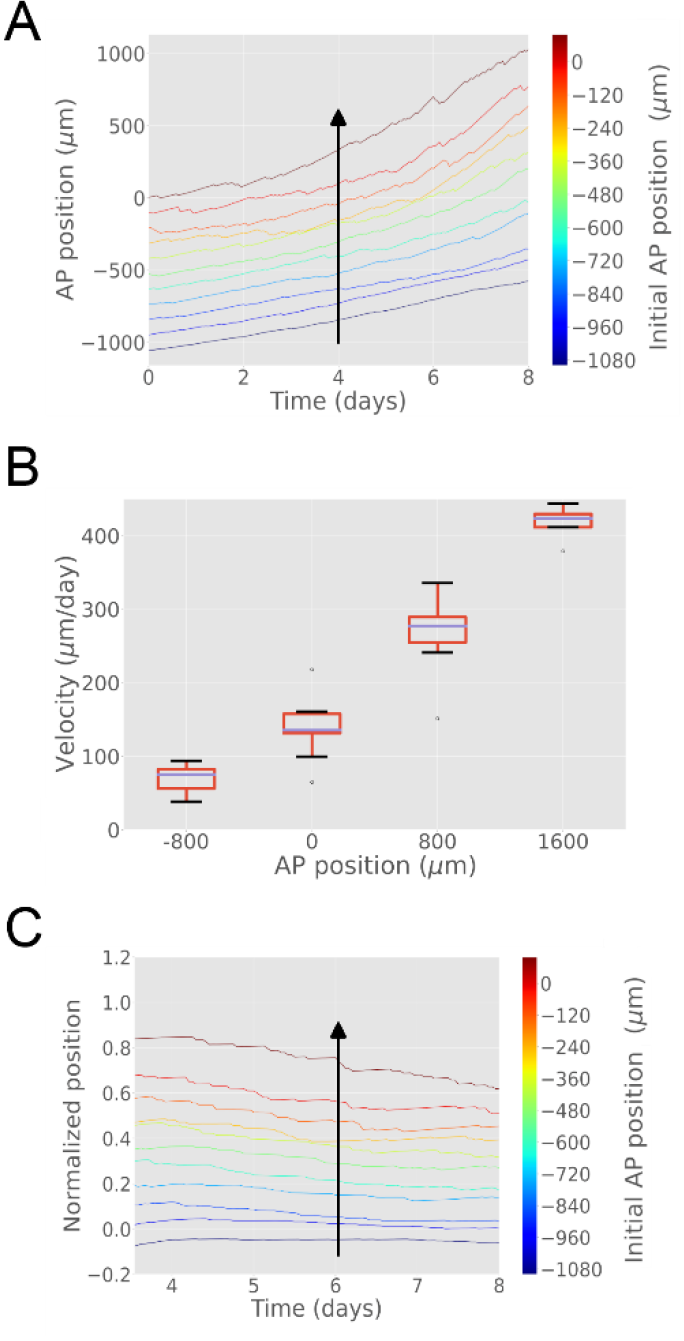
The model encompasses the cell pushing mechanism: the more posterior a cell is, the faster it moves. **A) Cells located close to the amputation plane end up at the posterior end of the regenerated modelled tissue. B) Clones velocity monotonically increases with AP coordinate.** Box plot representation showing median of clone velocities binned every 800 μm **C) Scaling behavior: clone cells preserve their original spatial order.** Relative position of each clone to the highly proliferating tissue delimited by its outgrowth and the recruitment limit remains constant in time. The figure depicts 11 simulations.

**Figure 2-figure supplement 3.**
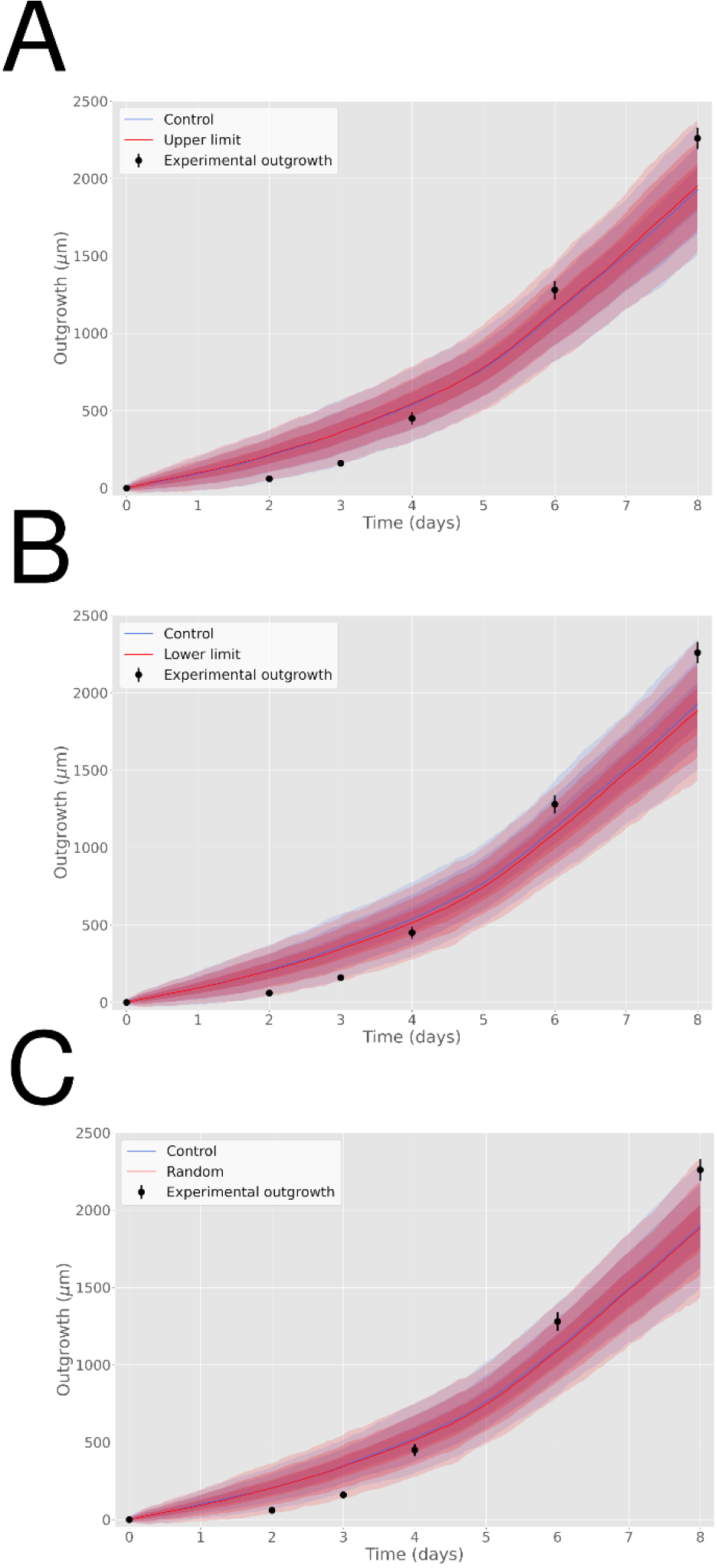
Incorporating variability of cell length along the AP axis does not impact on predicted axolotl spinal cord outgrowth. **(A)** Outgrowth predicted from the model assuming ependymal cell lengths equal to the experimental mean value plus three sigma of their cell length measured along the AP axis (in red). **(B)** Outgrowth predicted from the model assuming ependymal cell lengths equal to the experimental mean value minus three sigma of their cell length measured along the AP axis (in red). **(C)** Outgrowth predicted from the model assuming ependymal cell lengths extracted from a normal distribution parametrized from the experimental cell length measured along the AP axis (in red). In the three panels it is depicted the outgrowth predicted from the model assuming ependymal cell lengths equal to the experimental mean value of their cell length measured along the AP axis (Control, in blue, same simulations shown in Figure 2B). Experimental ependymal cell length along the AP axis are characterized by *μ* = 13.2 μm, *σ* = 0.1 μm.

**Figure 2-figure supplement 4.**
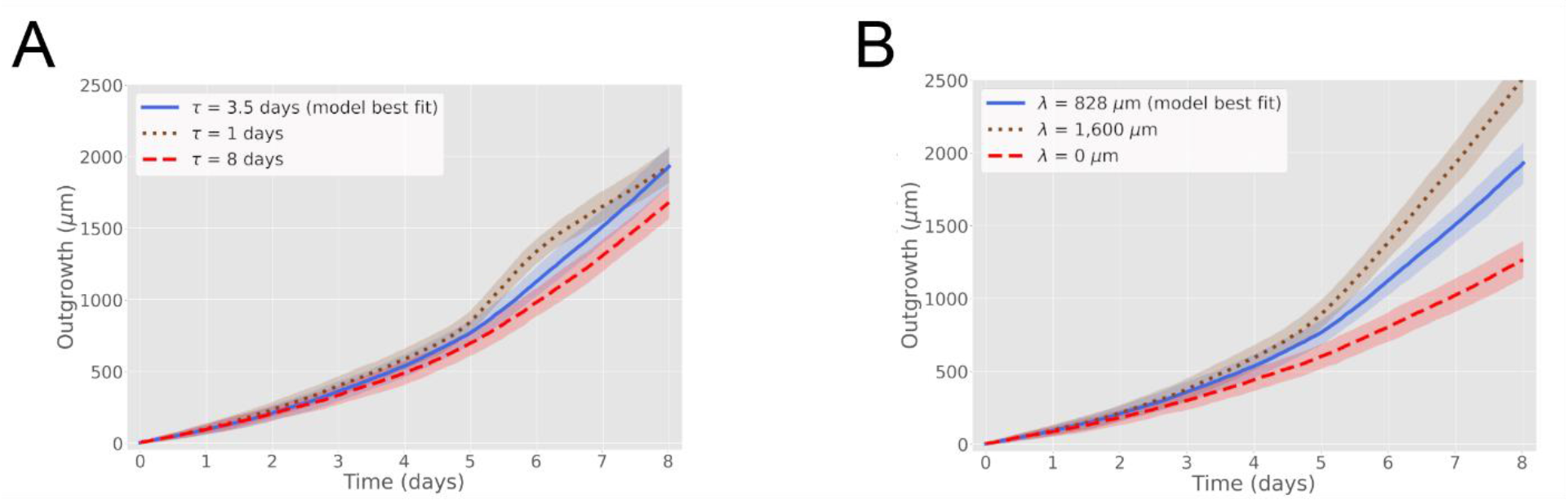
The maximal recruitment time (*τ*) and the maximal recruitment length (*λ*) determine the outgrowth of the axolotl spinal cord. **A) Increase (decrease) of *τ* reduces (increases) the model-predicted spinal cord outgrowth.** Time course of spinal cord outgrowth predicted by the model when varying *τ* from 1 day (brown) to 8 days (red). In blue, the prediction of the model assuming *τ* = 85 ± 12 hours. *N_0_* = 196 ± 2 cells and *λ* = 830 ± 30 μm. **B) Increase (decrease) of *λ* increases (reduces) the model-predicted spinal cord outgrowth.** Time course of spinal cord outgrowth predicted by the model when varying *λ* from zero (red) to 1,600 μm (brown). In blue, the prediction of the model assuming *λ* = 828 ± 30 μm. *N_0_* = 196 ± 2 cells and *τ* = 85 ± 12 hours. Simulations depicted in the blue curves in A and B are the same simulations shown in Fig. 2B. Means are depicted as lines while shaded areas correspond to 68 % confidence intervals, respectively, calculated from 1000 simulations.

**Figure 3-figure supplement 1.**
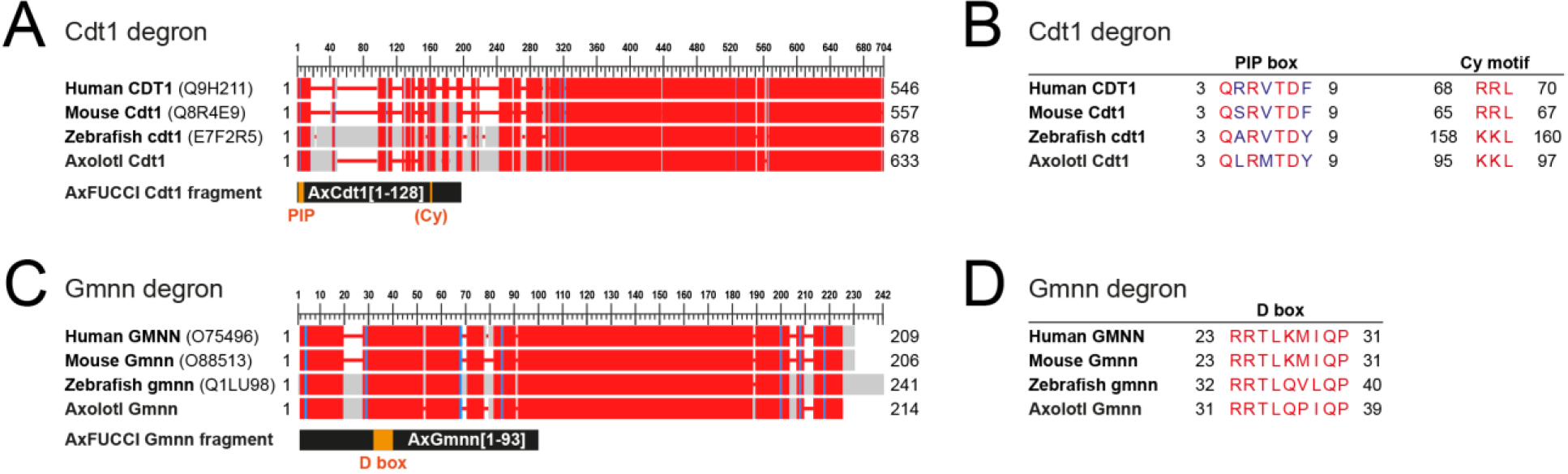
Determination of Cdt1 and Gmnn fragments for AxFUCCI reporter. **A) Alignment of full length Cdt1 proteins from human, mouse, zebrafish and axolotl.** Uniprot identifiers are given in parentheses. Numbers indicate amino acid residue numbers. Red regions indicate high amino acid similarity/identity; grey regions indicate low conservation. Alignments were performed using COBALT (NCBI). The AxFUCCI Cdt1 fragment comprises amino acids 1-128 of axolotl Cdt1, harboring the PIP box degron and corresponding to amino acids 1-190 of zebrafish cdt1 used to make zebrafish FUCCI (Sugiyama *et al*., 2009, Bouldin *et al*., 2014). **B) Amino acid alignments at the Cdt1 degron sequences.** The PIP box is well conserved evolutionarily. Like zebrafish cdt1, axolotl Cdt1 harbors a ‘KKL’ motif rather than the ‘RRL’ motif found in human and mouse. **C) Alignment of full length Gmnn proteins from human, mouse, zebrafish and axolotl.** The AxFUCCI Gmnn fragment comprises amino acids 1-93 of axolotl Gmnn, harboring the Destruction (D) box and corresponding to amino acids 1-100 of zebrafish gmnn1 used to make zebrafish FUCCI (Sugiyama *et al*., 2009, Bouldin *et al*., 2014). **D) Amino acid alignments at the Gmnn D box.**

**Figure 3-figure supplement 2.**
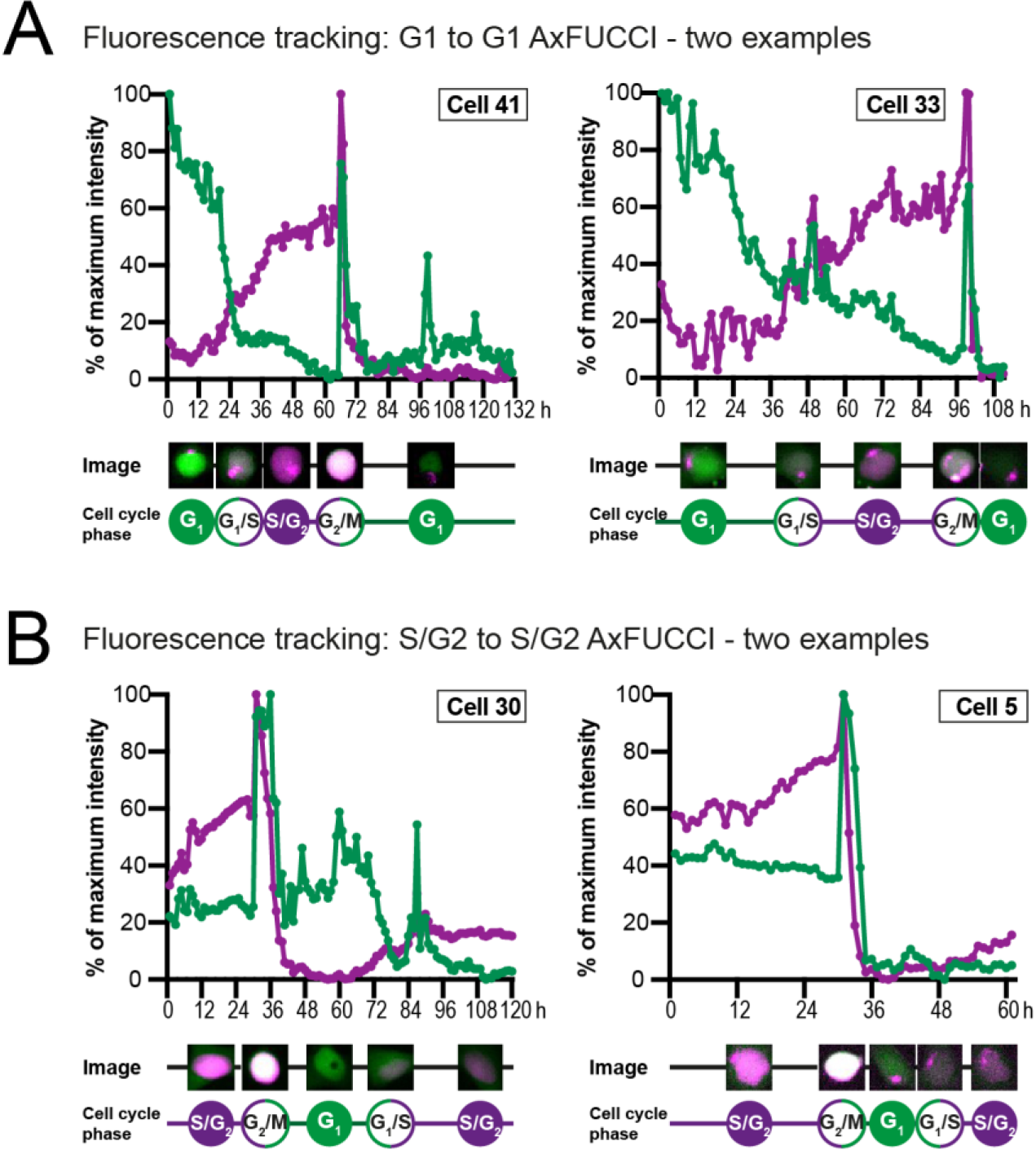
Fluorescence intensity measurements of AxFUCCI through the cell cycle. Fluorescence intensity measurements of AxFUCCI-electroporated AL1 cells as they transited through one cell cycle from G1 to the following G1 (**A**) or S/G2 to the following S/G2 (**B**). Two examples are presented for each transition. Green lines represent G0/G1-AxFUCCI fluorescence; magenta lines represent S/G2-AxFUCCI fluorescence. For each time point, the cell nucleus was segmented and the mean fluorescence intensity calculated. Fluorescence intensities were normalized to the maximal fluorescence value observed during the imaging session. h: hours after start of imaging. Maximum fluorescence is observed at mitosis, when the cell rounds up in preparation for cell division. Mean fluorescence intensity decreases sharply after each mitosis due to fluorophore and plasmid dilution between the two daughter cells. For sample videos, see Videos 2 and 3.

**Figure 3-figure supplement 3.**
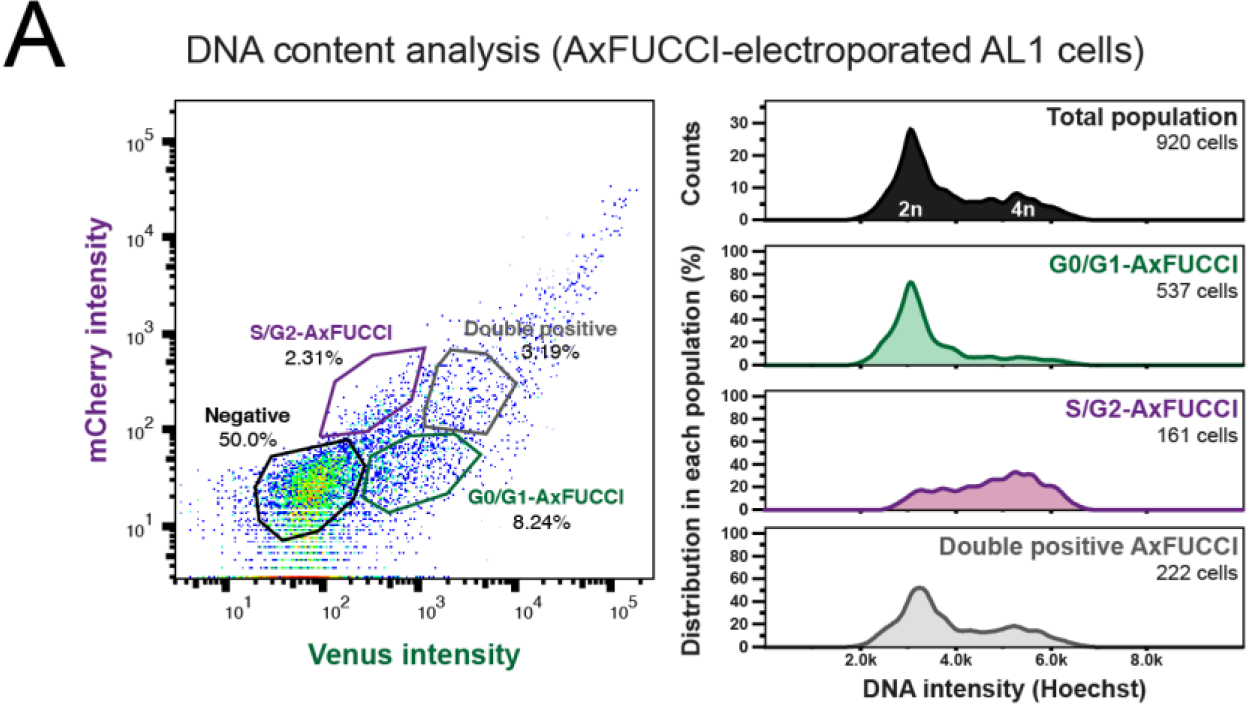
A) DNA quantification by flow cytometry to validate AxFUCCI functionality. **A)** Relative DNA content in G0/G1-AxFUCCI (green), S/G2-AxFUCCI (magenta) or Transition-AxFUCCI (grey) cells. AxFUCCI-electroporated AL1 cells were incubated with Hoechst DNA stain-containing cell culture medium for 90 minutes prior to dissociation and analysis.

**Figure 3-figure supplement 4.**
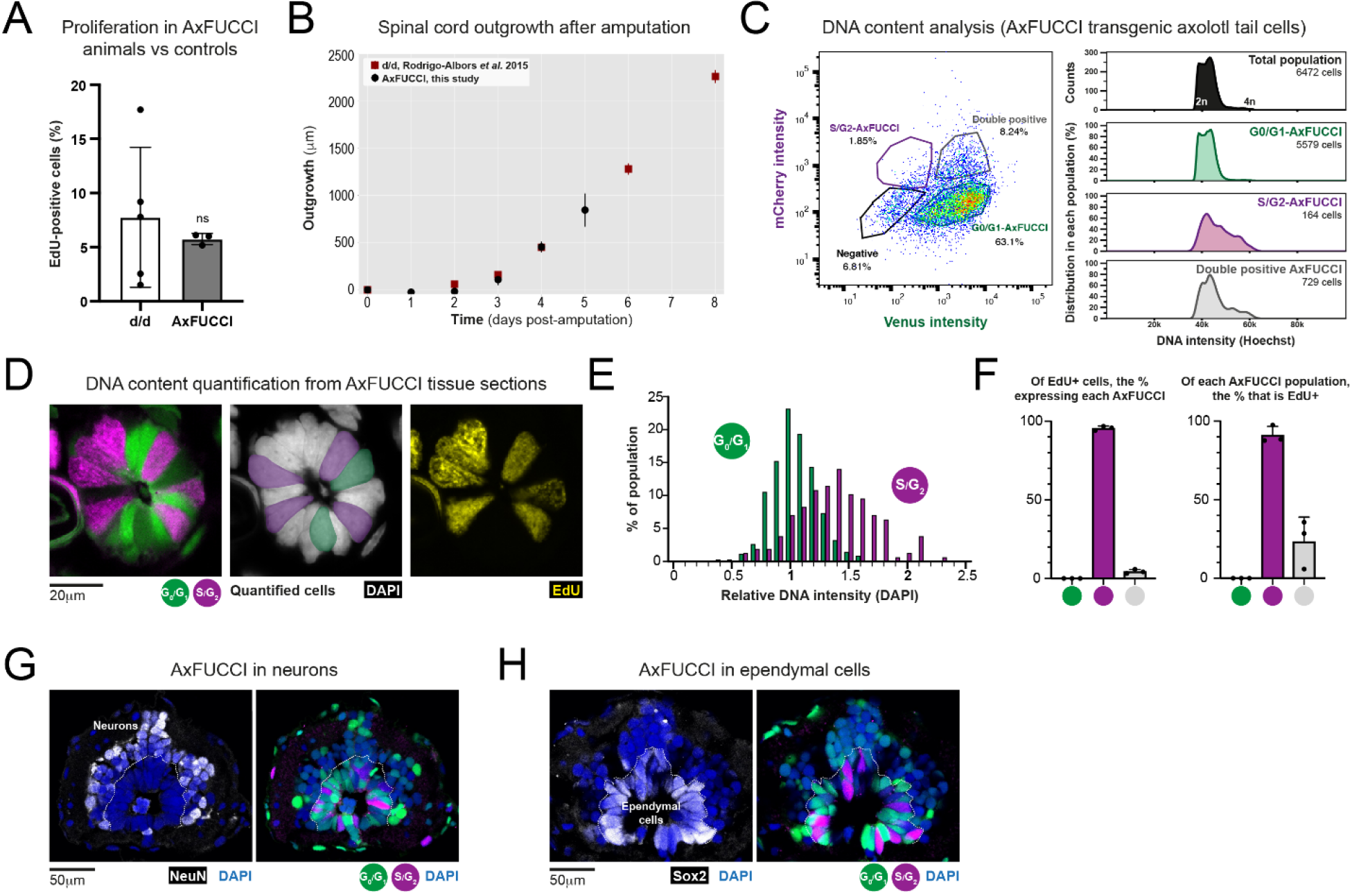
Validation of AxFUCCI functionality through *in vivo* tissue analysis. **A) Baseline proliferation in AxFUCCI spinal cord is not significantly different from d/d controls.** EdU was administered in a 2 hour pulse to uninjured AxFUCCI or d/d control animals. Tails were harvested and sectioned to quantify EdU-labelled proliferative cells in the spinal cord. ns: not significant, *p*=0.62 (two sample *t*-test). Error bars indicate standard deviation. *n=5* tails (control) or 3 tails (AxFUCCI), 750 cells counted in total for each, of which ~50 cells were EdU-labelled. **B) AxFUCCI tails regenerate with normal kinetics.** Spinal cord outgrowth was measured every day up to 5 days after AxFUCCI tail amputation, and plotted together with measurements previously obtained from *d/d* control animals (Rodrigo-Albors *et al*., 2015). **C) DNA quantification by flow cytometry.** Relative DNA content in G0/G1-AxFUCCI (green), S/G2-AxFUCCI (magenta) or Transition-AxFUCCI (grey) cells dissociated and harvested from uninjured AxFUCCI axolotl tails. Cells were incubated with Hoechst DNA stain for 60 minutes prior to analysis. **D) DNA quantification from AxFUCCI tissue sections.** Single section confocal image of a 10μm thick AxFUCCI spinal cord cross-section, co-stained with DAPI (grey) to label DNA. The DNA intensity of G0/G1-AxFUCCI cells (green) was compared to that of S/G2-AxFUCCI cells (magenta). The center panel depicts the cells that satisfy the criteria for quantification – they must span the spinal cord, contact the lumen and not be obscured by neighbouring cells. The AxFUCCI animals were also pulsed with EdU for 8 hours prior to tail harvesting. S/G2-AxFUCCI cells, but not G0/G1-AxFUCCI cells, incorporated EdU (yellow, right panel). **E) Histogram depicting the DAPI intensities of AxFUCCI cells quantified in D.** The mean intensity of G0/G1-AxFUCCI cells was set to an arbitrary value of 1 (2*n*). S/G2-AxFUCCI cells have a significantly higher DNA content than G0/G1-AxFUCCI cells (*p*=2.20×10^-16^, Welch’s test). *n*=3 tails (a total of 341 G0/G1-AxFUCCI cells vs 157 S/G2-AxFUCCI cells). **F) Correlation of EdU labelling (see panel D) with each AxFUCCI-expressing population.** *n*=3 tails, 819 AxFUCCI-expressing cells were counted in total, of which 171 had incorporated EdU. **G) Spinal cord neurons express G0/G1-AxFUCCI.** NeuN-expressing neurons are located on the periphery of the spinal cord. 98±1.5% of neurons expressed G0/G1-AxFUCCI, and the remainder did not express any AxFUCCI. *n*=5 tails, a total of 2,170 NeuN+ cells counted. **H) Ependymal cells express all AxFUCCI combinations.** Sox2-expressing ependymal cells are located on the interior of the spinal cord, lining the lumen.

**Figure 3-figure supplement 5.**
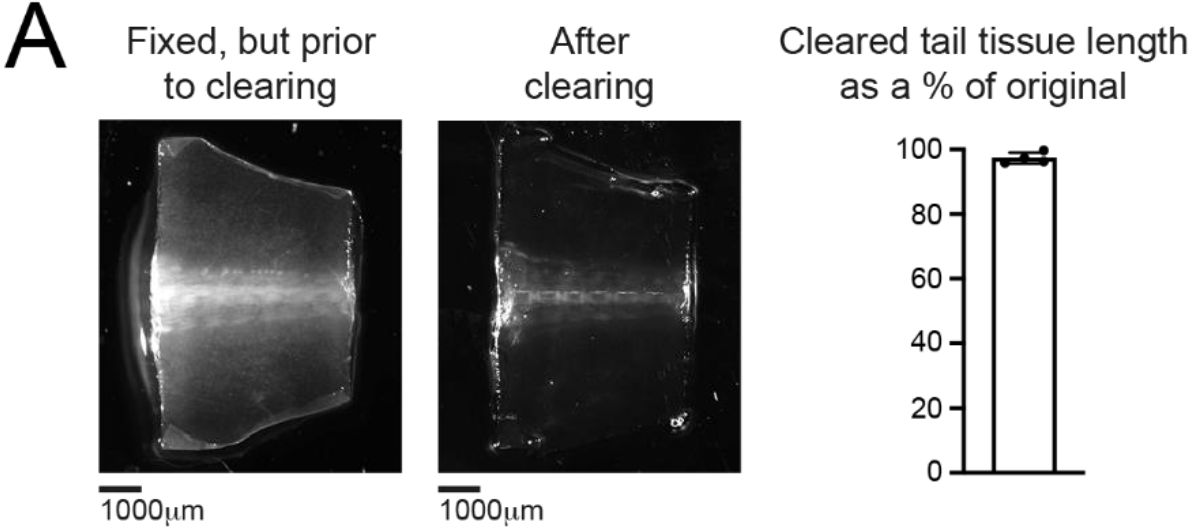
The tissue clearing protocol does not alter spinal cord length. **A)** The same, fixed tail tissue imaged prior to (left) or following (center) tissue clearing and refractive index matching. Right: Cleared tail tissue preserves 97±1.7% of its length compared to uncleared tail tissue. *n*=3 tails.

**Figure 4-figure supplement 1.**
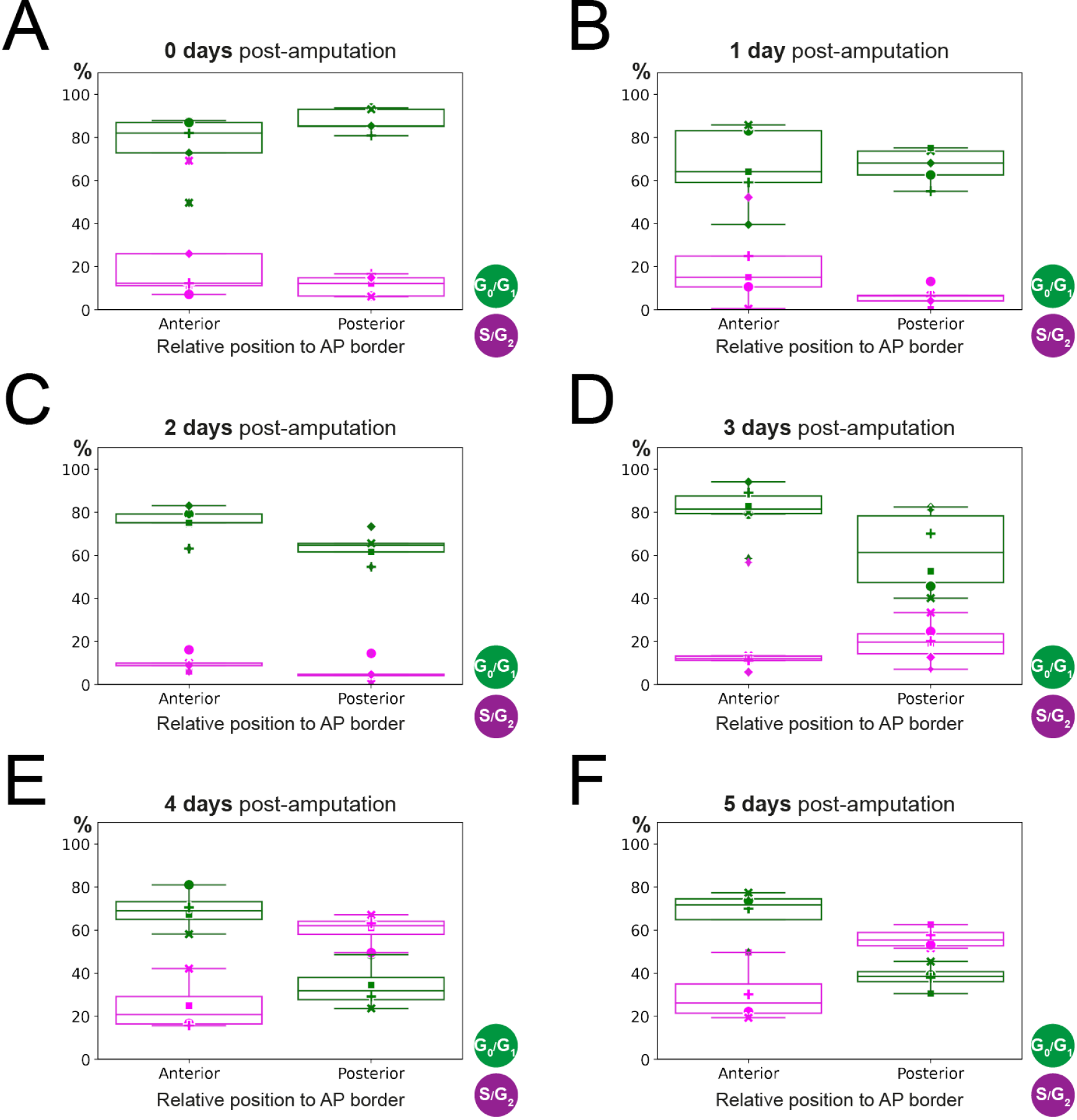
Estimation of anterior and posterior percentages of G0/G1-AxFUCCI and S/G2-AxFUCCI expressing cells by fitting a two-zones model. Dots correspond to the best-fitting value of the parameters *g*0*g*1_*a*_, *g*0*g*1_*p*_, *sg*2_*a*_ and *sg*2_*p*_ to the experimental spatial AP profiles of the percentages of G0/G1-AxFUCCI (in green) and S/G2-AxFUCCI (in magenta) expressing cells for each animal, at each timepoint **(A-E)**. Each symbol identifies a different animal (same criteria as in Figure 4). Box plots show that at day 0 to 3 days post-amputation, anterior and posterior values are not significantly different from each other (A – D). In contrast, at day 4 and 5, anterior and posterior values are significantly different (E, F) (Kolmogorov-Smirnov test *p* = 0.0286) (Materials and method section 2.12 for more details).

**Figure 4-figure supplement 2.**
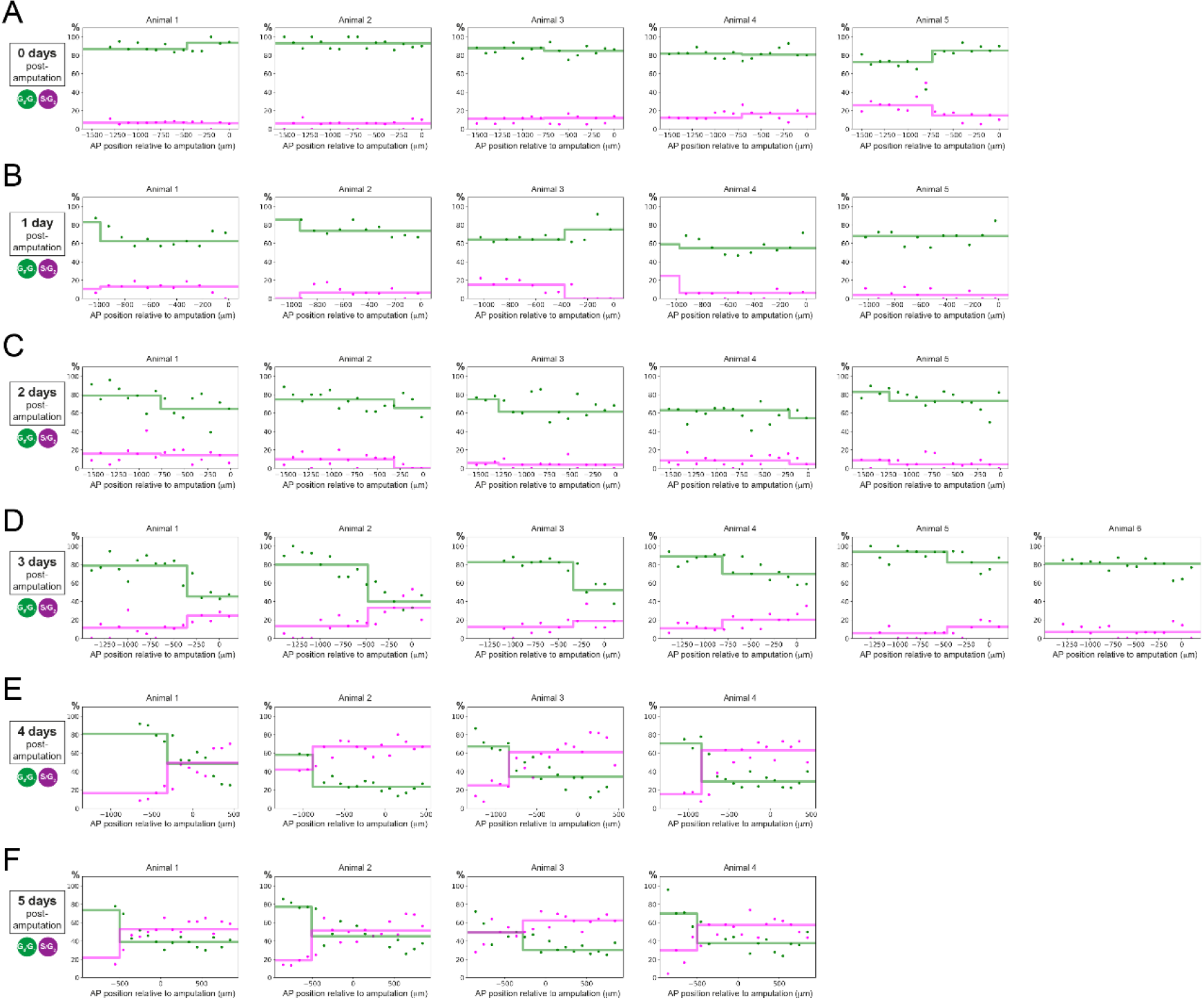
Individual fitting of the two-zones model to experimental AP profiles of the percentages of G0/G1-AxFUCCI and S/G2-AxFUCCI expressing cells. Percentage of G0/G1 (in green) and S/G2 (in magenta)-AxFUCCI-expressing cells quantified in 100 μm bins along the anterior-posterior spinal cord axis. A mathematical model assuming two adjacent spatially homogeneous zones separated by an anterior-posterior border (AP border) was fitted to the G0/G1 and S/G2-AxFUCCI-data for each animal at time 0 **(A)**, 1 **(B)**, 2 **(C)**, 3 **(D)**, 4 **(E)** and 5 **(F)** days post amputation. Anterior-posterior position is defined with respect to the amputation plane (0 μm). Best fitting values of the model regarding the anterior and posterior percentage of AxFUCCI data is in Figure 4 – figure supplement 1 (For more details, see Materials and Methods section 2.12).

**Figure 5-figure supplement 1.**
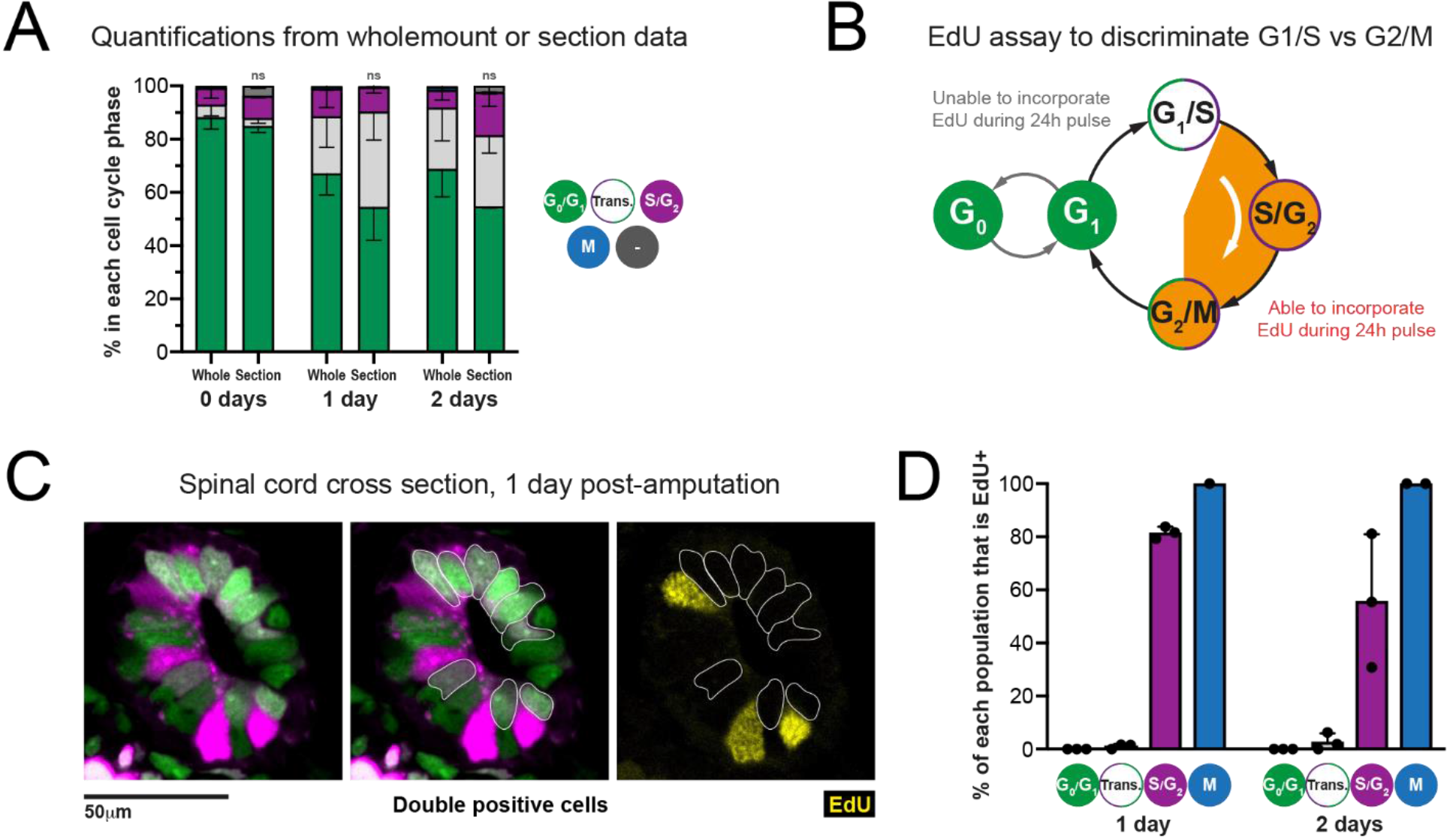
Transition-AxFUCCI cells at 1 and 2 days post-amputation reside at the G1/S boundary. **A) Comparison of AxFUCCI quantifications made from wholemount specimens (Whole) or tissue sections (Section).** No difference was observed between wholemount-based quantifications and section-based quantifications (one way ANOVA). *n*=3 tails per time point, ~400 cells counted in each, corresponding to ~750μm of spinal cord. Error bars indicate standard deviation. **B) EdU-based assay to distinguish G1/S cells and G2/M cells. C) Single 10μm section confocal image of a spinal cord at 1 day post-amputation, pulsed for 24 hours with EdU prior to harvesting.** Transition-AxFUCCI cells (double positive, outlined) have not incorporated EdU (yellow). Neighbouring S/G2-AxFUCCI cells (magenta) have incorporated EdU (internal control). **D) The percentage of each AxFUCCI population and mitotic cells that incorporated EdU in the assay in B.** As expected, no G0/G1-AxFUCCI cells incorporated EdU and all mitotic cells had incorporated EdU. The majority of S/G2-AxFUCCI cells had incorporated EdU. In contrast, almost none of the Transition-AxFUCCI cells at 1 and 2 days post-amputation incorporated EdU, indicating that they reside at the G1/S transition. *n*=3 tails per time point. Total cells counted per time point: 1 day post-amputation [671 G0/G1-AxFUCCI, 115 S/G2-AxFUCCI, 442 Transition-AxFUCCI, 1 mitotic cell], 2 days post-amputation [643 G0/G1-AxFUCCI, 191 S/G2-AxFUCCI, 315 Transition-AxFUCCI, 3 mitotic cells].

**Figure 6 – figure supplement 1.**
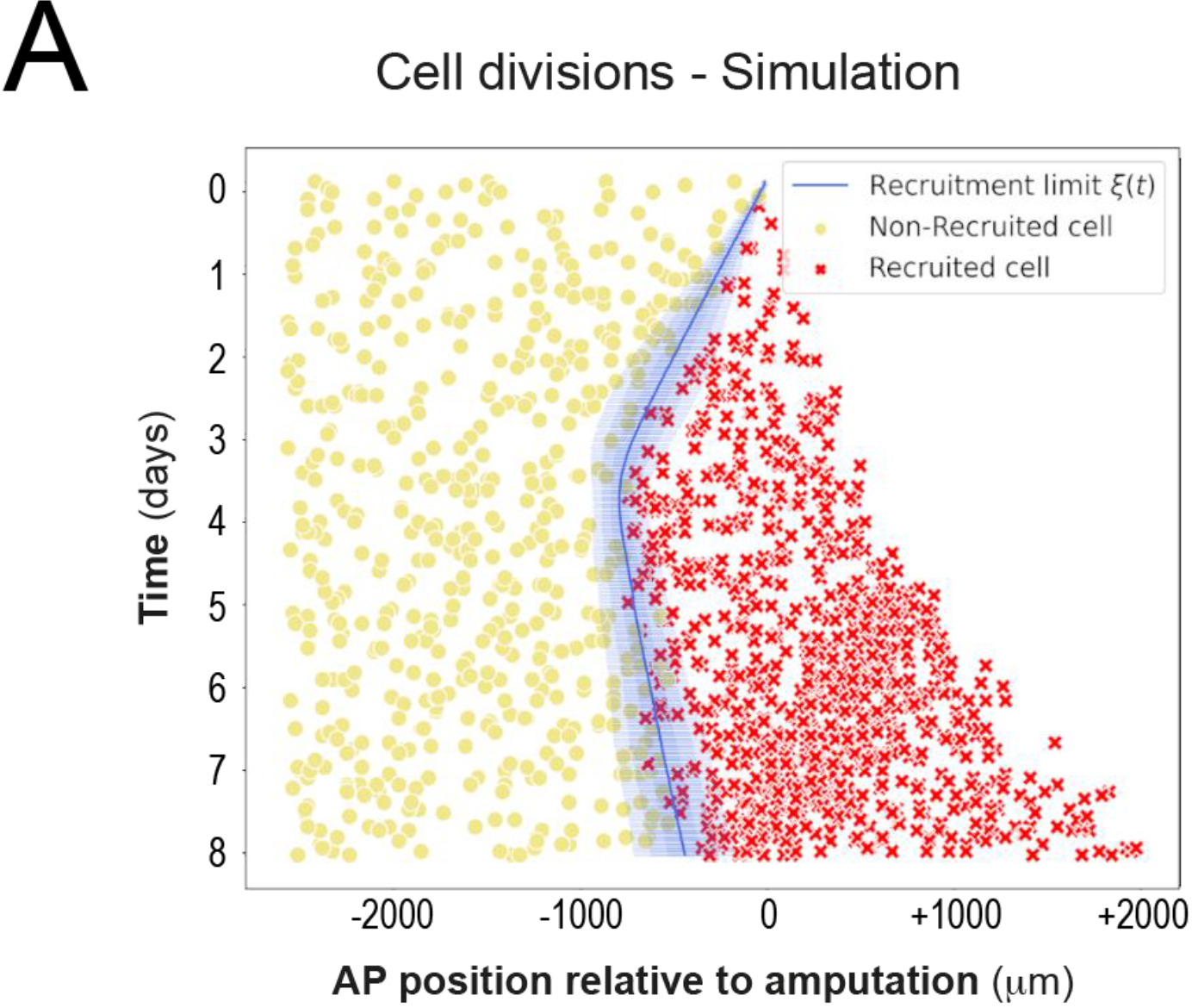
Modelled spatiotemporal distributions of ependymal cell divisions during axolotl spinal cord regeneration. Model-predicted occurrence of cell divisions is more often within the recruited (red crosses) compared to the non-recruited (yellow circles) cells, where recruited and non-recruited cells are separated by the recruitment limit *ξ*(*t*). The model is parameterized as in Figure 2 and Figure 6 C-E.

**Figure 6 – figure supplement 2.**
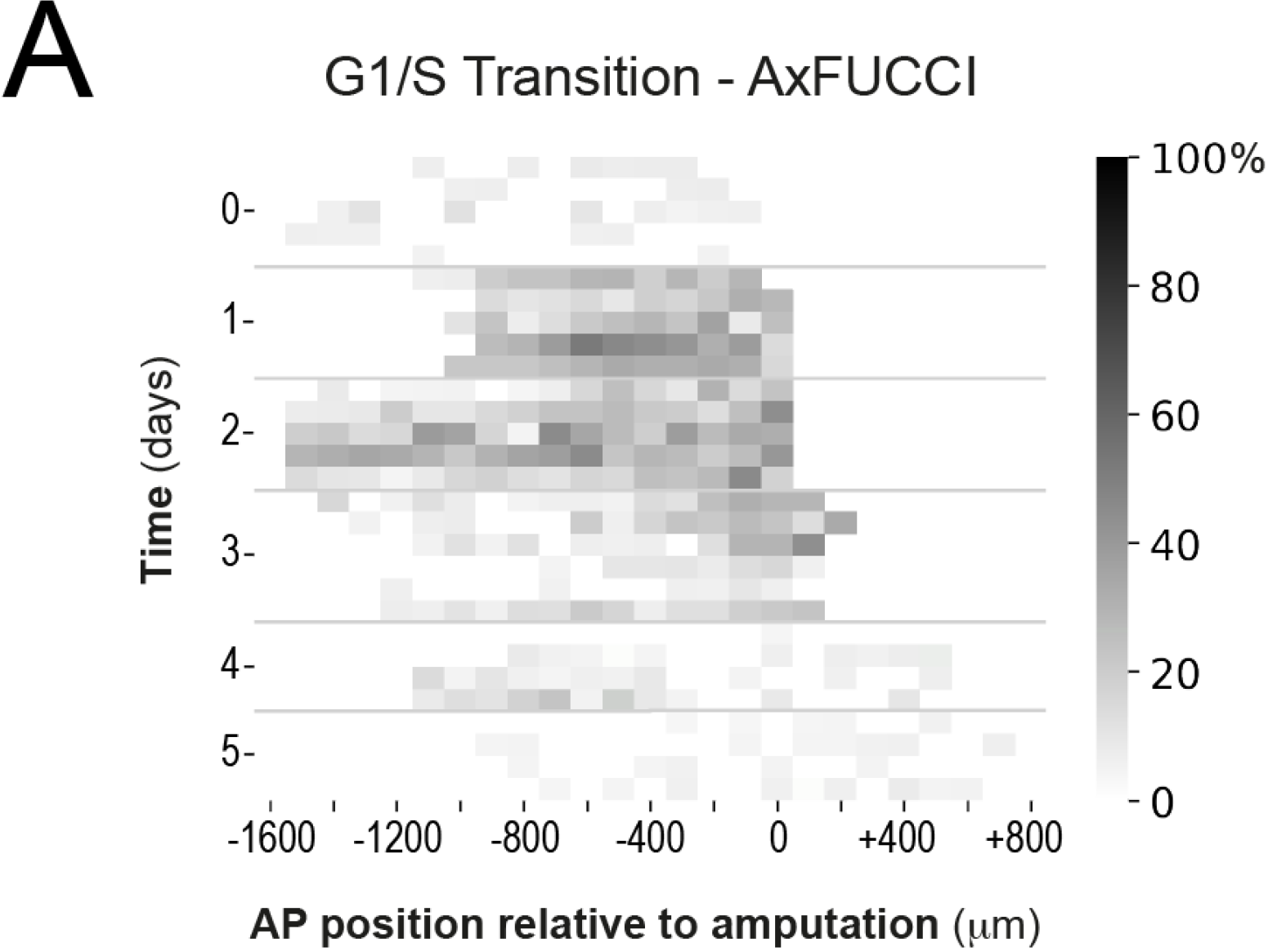
Spatiotemporal distributions of Transition AxFUCCI cells during axolotl spinal cord regeneration. Heatmaps depicting experimental spatiotemporal distribution of Transition AxFUCCI cells. Anterior-Posterior (AP) position is depicted horizontally with respect to the amputation plane (AP position = 0 μm). In the vertical axis time from amputation (Time = 0 days) and experimental replicates within each time. The color code corresponds to the percentage of Transition AxFUCCI cells. The increase in Transition AxFUCCI cells at days 1 and 2 post-amputation is readily observed.

## Video legends

**Video 1. 1D Model simulations of spinal cord regeneration. Top panel)** 20 model simulations from 20 different random seeds (the seeds are shown in the vertical axis). The color code corresponds to the generation of each cell (blue, orange and black correspond to the first, second and third generation, respectively). Vertical interrupted black line denotes the amputation plane (AP coordinate of 0). The recruitment limit *ξ*(*t*) (vertical red dashed line) propagates linearly in time up until the time *τ* of 85 hours post amputation, covering up the maximal recruitment *λ* of 828 μm anterior to the amputation plane. **Bottom left panel)** Predicted recruitment limit *ξ*(*t*) as a function of time from the simulations showed in the top panel). Mean value is depicted in the red line while the red shaded areas corresponds to 68, 95 and 99.7 % confidence intervals **Bottom right panel)** Predicted spinal cord outgrowth predicted by the model from the simulations showed in the top panel). The line represents to the mean (also indicated as the blue vertical line in the top panel) and the blue shaded areas correspond to the 68, 95 and 99.7 % confidence intervals. The 20 simulations have the same parametrization than the 1,000 simulations showed in Figure 2A and Figure 2B.

**Video 2. Live imaging of an AxFUCCI-electroporated AL1 cell (G1 to G1).** A single AxFUCCI-electroporated AL1 cell passing through one cell cycle from G1 to G1, imaged hourly over ~130h. The cell transitions through the cell cycle phases in the following order: Green (G0/G1-AxFUCCI) > White (Transition-AxFUCCI) > Magenta (S/G2-AxFUCCI) > White (Transition-AxFUCCI) > Mitosis > Two Green daughter cells (G0/G1-AxFUCCI). This cell corresponds to Cell 41 tracked in Figure 3 – figure supplement 2. Fluorescence intensity drops after mitosis due to fluorophore and plasmid dilution (non-integrating transgene).

**Video 3. Live imaging of an AxFUCCI-electroporated AL1 cell (S/G2 to S/G2).** A single AxFUCCI-electroporated AL1 cell passing through one cell cycle from S/G2 to S/G2, imaged overly over ~120h. The cell transitions through the cell cycle phases in the following order: Magenta (S/G2-AxFUCCI) > White (Transition-AxFUCCI) > Mitosis > Two Green daughter cells (G0/G1-AxFUCCI) > White (Transition-AxFUCCI) > Magenta (S/G2-AxFUCCI). This cell corresponds to Cell 30 tracked in Figure 3 – figure supplement 2. Fluorescence intensity drops after mitosis due to fluorophore and plasmid dilution (non-integrating transgene).

**Video 4. 3D imaging of an optically cleared AxFUCCI tail tip.** Volume rendering of a fixed and optically cleared AxFUCCI tail tip, co-stained for DNA using DAPI (blue) and imaged with a lightsheet microscope. The spinal cord is visible as an intensely green and magenta rod at the center of the sample. Peripheral signal is AxFUCCI expression in surface epidermal cells. Other internal structures, including notochord, have weaker AxFUCCI expression and are not visible in this rendering. Volume rendering was performed using Imaris software. See also Figure 3D.

**Video 5. 1D Model simulations of spinal cord regeneration showing cells in G1 and S phases**. 20 model simulations from 20 different random seeds (the seeds are shown in the vertical axis). Cells in G1 and S are depicted in green and magenta, respectively. Predicted recruitment limit *ξ*(*t*) and outgrowth are indicated with the vertical red and blue discontinuous lines, respectively. Amputation plane is depicted as a vertical black dashed line. The simulations have the same parametrization than the simulations showed in Figure 2A and Figure 2B.

## References

Abe T, Sakaue-Sawano A, Kiyonari H, Shioi G, Inoue K, Horiuchi T, Nakao K, Miyawaki A, Aizawa S & Fujimori T. 2013. Visualization of cell cycle in mouse embryos with Fucci2 reporter directed by Rosa26 promoter. Development. 140:237–246.

Adams DS, Masi A & Levin M. 2007. H+ Pump-dependent changes in membrane voltage are an early mechanism necessary and sufficient to induce Xenopus tail regeneration. Development. 34(7): 1323–35.

Arai Y, Pulvers JN, Haffner C, Schilling B, Nüsslein I, Calegari F, Huttner WB. 2011. Neural stem and progenitor cells shorten S-phase on commitment to neuron production. Nat Commun. 2:154.

Becker CG, Becker T, Hugnot JP. 2018. The spinal ependymal zone as a source of endogenous repair cells across vertebrates. Prog Neurobiol. 170: 67–80.

Bouldin CM, Snelson CD, Farr GH 3^rd^ & Kimelman D. 2014. Restricted expression of cdc25a in the tailbud is essential for formation of the zebrafish posterior body. Genes Dev. 28:384–395.

Calegari F & Huttner WB. 2003. An inhibition of cyclin-dependent kinases that lengthens, but does not arrest, neuroepithelial cell cycle induces premature neurogenesis. J Cell Sci. 116: 4947–4955.

Calegari F, Haubensak W, Haffner C, Huttner WB. 2005. Selective lengthening of the cell cycle in the neurogenic subpopulation of neural progenitor cells during mouse brain development. J Neurosci. 25(28): 6533–8.

Chara O, Tanaka EM & Brusch L. 2014. Mathematical modeling of regenerative processes. Curr Top Dev Biol. 108: 283–317.

Chernoff EA, Stocum DL, Nye HL, Cameron JA. 2003. Urodele spinal cord regeneration and related processes. Dev Dyn. 226(2): 295–307.

Chiou K & Collins ES. 2018. Why we need mechanics to understand animal regeneration. Dev Biol. 433(2): 155–165.

Csilléry K, Blum MGB, Gaggiotti OE & François O. 2010. Approximate Bayesian Computation (ABC) in practice. Trends in Ecology & Evolution. 25 (7): 410–418.

Cura Costa E, Otsuki L, Rodrigo Albors A, Tanaka EM & Chara O. 2021. Spatiotemporal control of cell cycle acceleration during axolotl spinal cord regeneration - Supplementary notebooks – v2.0. Zenodo. https://doi.org/10.5281/zenodo.4557840

Fei JF, Schuez M, Tazaki A, Taniguchi Y, Roensch K & Tanaka EM. 2014. CRISPR-mediated genomic deletion of Sox2 in the axolotl shows a requirement in spinal cord neural stem cell amplification during tail regeneration. Stem Cell Reports. 3(3): 444–59.

Fei JF, Knapp D, Schuez M, Murawala P, Zou Y, Singh SP, Drechsel D & Tanaka EM. 2016. Tissue- and time-directed electroporation of CAS9 protein–gRNA complexes in vivo yields efficient multigene knockout for studying gene function in regeneration. npj Regen Med. 1(1): 16002.

Freitas PD, Yandulskaya AS & Monaghan JR. 2019. Spinal Cord Regeneration in Amphibians: A Historical Perspective. Dev Neurobiol. 79(5): 437–452.

Hasenpusch-Theil K, West S, Kelman A, Kozic Z, Horrocks S, McMahon AP, Price DJ, Mason JO, Theil T. 2018. Gli3 controls the onset of cortical neurogenesis by regulating the radial glial cell cycle through Cdk6 expression. Development. 145(17).

Harris CR, Jarrod Millman K, van der Walt SJ, Gommers R, Virtanen P, Cournapeau D, Wieser E, Taylor J, Berg S, Smith NJ, Kern R, Picus M, Hoyer S, van Kerkwijk MH, Brett M, Haldane A, Fernández del Río J, Wiebe M, Peterson P, Gérard-Marchant P, Sheppard K, Reddy T, Weckesser W, Abbasi H, Gohlke C & Oliphant TE. 2020. Array programming with NumPy, Nature, 585, 357–362. DOI:10.1038/s41586-020-2649-2

Hunter JD. 2007. Matplotlib: A 2D graphics environment. Computing in Science & Engineering. 9:90–95.

Ilieş I, Sipahi R & Zupanc GKH. 2018. Growth of adult spinal cord in knifefish: development and parametrization of a distributed model. Journal of Theoretical Biology. 437: 101–114.

Joven A & Simon A. 2018. Homeostatic and regenerative neurogenesis in salamanders. Prog Neurobiol. 170: 81–98.

Kicheva A, Bollenbach T, Ribeiro A, Valle HP, Lovell-Badge R, Episkopou V & Briscoe J. 2014. Coordination of progenitor specification and growth in mouse and chick spinal cord. Science 345: 1254927.

Klinger E, Rickert D, Hasenauer J. 2018. pyABC: distributed, likelihood-free inference. Bioinformatics. 34(20): 3591–3593.

Lander AD, Gokoffski KK, Wan FY, Nie Q & Calof AL. 2009. Cell lineages and the logic of proliferative control. PLoS Biol. 7(1): e15.

Lange C, Huttner WB & Calegari F. 2009. Cdk4/cyclinD1 overexpression in neural stem cells shortens G1, delays neurogenesis, and promotes the generation and expansion of basal progenitors. Cell Stem Cell. 5(3): 320–31.

Lukaszewicz A, Savatier P, Cortay V, Giroud P, Huissoud C, Berland M, Kennedy H & Dehay C. 2005. G1 phase regulation, area-specific cell cycle control, and cytoarchitectonics in the primate cortex. Neuron 47: 353–364.

McKinney W. 2010. Data structures for statistical computing in Python. Proceedings of the 9th Python in Science Conference: 51–56.

Morelli LG, Uriu K, Ares S & Oates AC. 2012. Computational approaches to developmental patterning. Science. 336(6078): 187–91.

Nordman J & Orr-Weaver TL. 2012. Regulation of DNA replication during development. Development 139: 455–464.

Oliphant TE. 2006. A guide to NumPy, USA: Trelgol Publishing.

Ozkucur N, Epperlein HH & Funk RH. 2010. Ion imaging during axolotl tail regeneration in vivo. Dev Dyn. 239(7): 2048–57.

Pende M, Vadiwala K, Schmidbaur H, Stockinger AW, Murawala P, Saghafi S, Dekens MPS, Becker K, Revilla-i-Domingo R, Papadopoulos S-C, Zurl M, Pasierbek P, Simakov O, Tanaka EM, Raible F & Dodt HU. 2020. A versatile depigmentation, clearing, and labeling method for exploring nervous system diversity. Sci Adv. 6: eaba0365.

Pilaz LJ, Patti D, Marcy G, Ollier E, Pfister S, Douglas RJ, Betizeau M, Gautier E, Cortay V, Doerflinger N, Kennedy H & Dehay C. 2009. Forced G1-phase reduction alters mode of division, neuron number, and laminar phenotype in the cerebral cortex. Proc Natl Acad Sci U S A. 106(51): 21924–9.

Rodrigo Albors A, Tazaki A, Rost F, Nowoshilow S, Chara O & Tanaka EM. 2015. Planar cell polarity-mediated induction of neural stem cell expansion during axolotl spinal cord regeneration. eLife. 4: e10230.

Rost F, Rodrigo Albors A, Mazurov V, Brusch L, Deutsch A, Tanaka EM & Chara O. 2016. Accelerated cell divisions drive the outgrowth of the regenerating spinal cord in axolotls. eLife. 5. pii: e20357.

Sabin K, Santos-Ferreira T, Essig J, Rudasill S & Echeverri K. 2015. Dynamic membrane depolarization is an early regulator of ependymoglial cell response to spinal cord injury in axolotl. Dev Biol. 408(1): 14–25.

Sakaue-Sawano A, Kurokawa H, Morimura T, Hanyu A, Hama H, Osawa H, Kashiwagi S, Fukami K, Miyata T, Miyoshi H, Imamura T, Ogawa M, Masai H & Miyawaki A. 2008. Visualizing spatiotemporal dynamics of multicellular cell-cycle progression. Cell. 132:487–498.

Salomoni P & Calegari F. 2010. Cell cycle control of mammalian neural stem cells: putting a speed limit on G1. Trends Cell Biol. 20: 233–243.

Schindelin J, Arganda-Carreras I, Frise E, Kaynig V, Longair M, Pietzsch T, Preibisch S, Rueden C, Saalfeld S, Schmid B, Tinevez J-Y, White DJ, Hartenstein V, Eliceiri K, Tomancak P & Cardona A. 2012. Fiji: an open-source platform for biological-image analysis. Nature Methods. 9: 676–682.

Schlüßler R, Möllmert S, Abuhattum S, Cojoc G, Müller P, Kim K, Möckel C, Zimmermann C, Czarske J & Guck J. 2018. Mechanical Mapping of Spinal Cord Growth and Repair in Living Zebrafish Larvae by Brillouin Imaging. Biophys J. 115(5): 911–923.

Sobkow L, Epperlein HH, Herklotz S, Straube WL & Tanaka EM. 2006. A germline GFP transgenic axolotl and its use to track cell fate: dual origin of the fin mesenchyme during development and the fate of blood cells during regeneration. Dev. biol. 290:386–397.

Sugiura T, Wang H, Barsacchi R, Simon A & Tanaka EM. 2016. MARCKS-like protein is an initiating molecule in axolotl appendage regeneration. Nature. 531(7593): 237–40.

Sugiyama M, Sakaue-Sawano A, Iimura T, Fukami K, Kitaguchi T, Kawakami K, Okamoto H, Higashijima S & Miyawaki A. 2009. Illuminating cell-cycle progression in the developing zebrafish embryo. Proc Natl Acad Sci USA. 106:20812–20817.

Takahashi T, Nowakowski RS & Caviness VS Jr. 1995. The cell cycle of the pseudostratified ventricular epithelium of the embryonic murine cerebral wall. J Neurosci. 15(9): 6046–57.

Tazaki A, Tanaka EM & Fei JF. 2017. Salamander spinal cord regeneration: The ultimate positive control in vertebrate spinal cord regeneration. Dev Biol. 432(1): 63–71.

Tinevez JY, Perry N, Schindelin J, Hoopes GM, Reynolds GD, Laplantine E, Bednarek SY, Shorte SL & Eliceiri KW. 2017. TrackMate: An open and extensible platform for single-particle tracking. Methods. 115:80–90.

Toni T, Welch D, Strelkowa N, Ipsen A, Stumpf MP. 2009. Approximate Bayesian computation scheme for parameter inference and model selection in dynamical systems. J R Soc Interface. 6(31): 187–202.

Turrero García M, Chang Y, Arai Y & Huttner WB. 2016. S-phase duration is the main target of cell cycle regulation in neural progenitors of developing ferret neocortex. J Comp Neurol. 524(3): 456–70.

Virtanen P, Gommers R, Oliphant TE, Haberland M, Reddy T, Cournapeau D, Burovski E, Peterson P, Weckesser W, Bright J, van der Walt SJ, Brett M, Wilson J, Jarrod Millman K, Mayorov N, Nelson ARJ, Jones, Kern R, Larson E, Carey CJ, Polat Ì, Feng Y, Moore EW, VanderPlas J, Laxalde D, Perktold J, Cimrman R, Henriksen I, Quintero EA, Harris CR, Archibald AM, Ribeiro AH, Pedregosa F, van Mulbregt P, and SciPy 1.0 Contributors. 2020. SciPy 1.0: Fundamental Algorithms for Scientific Computing in Python. Nature Methods, 17(3), 261–272.

Waskom M, Botvinnik O, O’Kane D, Hobson P, Lukauskas S, Gemperline DC, Augspurger T, Halchenko Y, Cole JB, Warmenhoven J, de Ruiter J, Pye C, Hoyer S, Vanderplas J, Villalba S, Kunter G, Quintero E, Bachant P, Martin M, Meyer K, Miles A, Ram T, Yarkoni T, Lee Williams M, Evans C, Fitzgerald C, Brian, Fonnesbeck C, Lee A, Qalieh A. https://doi.org/10.5281/zenodo.883859

Zielke N & Edgar BA. 2015. FUCCI Sensors: powerful new tools for analysis of cell proliferation. WIREs Dev Biol. 4:469–487.

